# A transcriptome based aging clock near the theoretical limit of accuracy

**DOI:** 10.1101/2020.05.29.123430

**Authors:** David H. Meyer, Björn Schumacher

## Abstract

Aging clocks dissociate biological from chronological age. The estimation of biological age is important for identifying gerontogenes and assessing environmental, nutritional or therapeutic impacts on the aging process. Recently, methylation markers were shown to allow estimation of biological age based on age-dependent somatic epigenetic alterations. However, DNA methylation is absent in some species such as *Caenorhabditis elegans* and it remains unclear whether and how the epigenetic clocks affect gene expression. Aging clocks based on transcriptomes have suffered from considerable variation in the data and relatively low accuracy. Here, we devised an approach that uses temporal scaling and binarization of *C. elegans* transcriptomes to define a gene set that predicts biological age with an accuracy that is close to the theoretical limit. Our model accurately predicts the longevity effects of diverse strains, treatments and conditions. The involved genes support a role of specific transcription factors as well as innate immunity and neuronal signaling in the regulation of the aging process. We show that this transcriptome clock can also be applied to human age prediction with high accuracy. This transcriptome aging clock could therefore find wide application in genetic, environmental and therapeutic interventions in the aging process.

## Introduction

Aging is the driving factor for several diseases, the declining organ function and overall progressive loss of physiological integrity^1^. Aging biomarkers that predict the biological age of an organism are important for identifying genetic and environmental factors that influence the aging process and for accelerating studies examining potential rejuvenating treatments. Initial studies have shown evidence that methods predicting the biological age are indeed sensitive enough to detect the effect of geroprotective therapies^2–5^. Diverse studies tried to identify biomarkers and predict the age of individuals, ranging from proteomics, transcriptomics, the microbiome, frailty index assessments to neuroimaging and DNA methylation^6–16^. Several age predictors based on easy to obtain medical records or hematological data have shown some promising results but are still lacking in overall accuracy^17–21^.

### DNA Methylation clocks

Currently, the most common predictors are based on DNA methylation. An elastic net regression model based on whole blood-derived 71-methylation sites could predict a validation cohort with a correlation coefficient of 0.91 and a root-mean-square error (RMSE) of 4.9 years (for details on parameters reported from the literature, see methods). Genes with nearby age-associated methylation markers predicted the age of 488 public available whole blood gene expression samples with an RMSE of 7.22 years^22^. The first multi-tissue predictor of age comprising 51 distinct tissue and cell types utilized 353 CpG methylation sites and resulted in a correlation of 0.96 and a median absolute difference of 3.6 years^23^. Consequently, a variety of epigenetic aging clocks in humans^24,25^ with as few as 8 CpG sites^26^ and also other organisms^3,27^ were devised. Despite these advances there are still challenges and weaknesses^28^: A recent report showed that the improved prediction of chronological age from DNA methylation might limit its use as a biomarker of aging. A hypothetical perfect prediction of the chronological age would not give any information of the biological age of the organism. The deviation of the prediction from the chronological age (so called age acceleration residual) can therefore give insight into the probable mortality. The improvement of the chronological age predictor is thereby accompanied by a decreased association between mortality and the bias of prediction^29^. The same study found a potential cellular composition confounder effect in those epigenetic clocks. After correction for this confounder, no significant association between the age acceleration residual and mortality was found, suggesting that the biggest effect is driven by differences in the cellular composition and thereby might limit the usage of the DNA methylation marks itself as biomarkers.

### Transcriptomics clocks

The DNA methylation marks themselves might influence the transcriptional response^22,30,31^, but aging also affects the transcriptional network by altering the histone abundance^32^, histone modifications^33–37^ as well as the 3D organization of chromatin^38,39^. The difference in RNA molecule abundance, thereby, integrates a variety of regulation and influences resulting in a notable gene expression change during the lifespan of an organism^40–46^, aging-associated changes in transcriptional elongation^47^ and a systemic length-driven transcriptome imbalance^48^. A recent study identified six gene expression hallmarks of cellular aging across eukaryotes from yeast to humans^49^ and the suppression of the transcriptional drift has been shown to extend the lifespan of *C. elegans*^50^. These studies sparked interest in the identification of transcriptomic aging biomarkers, an RNA expression signature for age classification and the development of transcriptomic aging clocks.

Peters et al. extended previous classification approaches^51–55^ to a regression, which allows the computation of the predicted age and developed a transcriptional aging clock based on whole-blood microarray samples for half of the human genome and reported an r^2^ of up to 0.6, an average difference of 7.8 years and an association of the predicted age to blood pressure as well as smoking status^56^. Similarly, Mamoshina et al. build a transcriptomic aging clock of human muscle tissue. A deep feature selection model performed best with an r^2^ of 0.83 and a mean absolute error of 6.24 years^57^. Recently, Affymetrix samples of the cortex, the hippocampus and the cerebellum were used to train a deep learning predictor to an r^2^ of 0.91 and an RMSE of 7.76 years^58^.

However, microarray data have the drawbacks of a limited range of detection, high background levels and the detection of just a subset of the transcriptome. To overcome these limitations, a transcriptional age predictor based on human RNA-seq data from the GTEx project^59^ yielded Spearman correlation coefficients of up to 0.84, dependent on the tissue^60^. By applying an ensemble of linear discriminant analysis classifiers, a model with an r^2^ of 0.81, a mean absolute error of 7.7 years and a median absolute error of 4.0 years was obtained in a dataset derived from cell culture of healthy donors^61^. The same model also predicted an accelerated age in 10 patients with the premature aging disease Hutchinson-Gilford progeria syndrome (HGPS). The first across-tissue transcriptional age calculator using a LASSO regression showed that splicing events could predict with an overall accuracy of 71 %^62^, while an across-tissue prediction on gene expression data showed that an elastic net regression was the most accurate with an average Pearson correlation coefficient across tissues of 0.33 ^63^. Apart from mRNA sequencing, a study of human peripheral blood micro RNAs samples was able to predict the age with an r^2^ of 0.49 ^64^. Moreover, it has been shown that for lung tissue of mice transcriptional clocks are more accurate than epigenetic clocks^65^.

### Proteomics clocks

Proteomics has been used in encouraging recent studies on human aging clocks and biomarker discovery. The first proteomics aging clock based on human blood plasma identified age-associated proteins that were used to build an elastic net prediction with an r^2^ of 0.88 ^66^, which could later be improved to an r^2^ of 0.94 ^67^. Recently, a study based on blood proteins predicted human age with a Pearson correlation of 0.88 more accurately than a combination of these proteins with metabolites and several clinical lab tests showing the versatility of also only a subset of proteins^68^. A review of 32 different human proteomics and aging studies aiming to identify common age-associated proteins that could be robustly identified regardless of the technique or population diversity used, resulted in an age predictor based on 83 proteins that were reported in three or more out of the 32 different studies and reported a Spearman correlation of 0.91 ^69^. This study also showed the current limitation of the usage of proteomics for age prediction: there is no standard in the generation of proteomic data and different techniques detect different subsets of proteins, which might lead to a measurement bias, i.e. some important age-related proteins might have been overlooked, while other were reported too frequently.

Summarizing, a large variety of data, techniques and analyses have been used to identify aging-biomarkers and -clocks in humans. However, these analyses also showed a pronounced variability and difficulties in replicability. Indeed, a recent analysis^70^ of gene expression, plasma protein, blood metabolite, blood cytokine, microbiome and clinical marker data^71,72^, showed that individual age slopes diverged among the participants over the longitudinal measurement time and subsequently that individuals have different molecular aging pattern, called ageotypes^73^. Moreover, a recent 10-year longitudinal study^74^ showed that individuals are more similar to themselves than to others with the same age and a twin study^75^ demonstrated that the global effect of age in gene expression is small. These interindividual differences are even more pronounced between different ethnicities and sex^19,76^ and show that it is still difficult to pinpoint biomarkers for aging in humans.

### *C. elegans* aging clocks

Model organisms, instead, can give a more controllable view on the aging process and biomarker discovery and several studies have been conducted in mice and rats^3,27,65,77–79^ and similarities between model organism and human aging have been described^80–82^. *C. elegans* has revolutionized the aging field and has vast advantages as a model organism^83–87^. Even isogenic nematodes in precisely controlled homogenous environments have surprisingly diverse lifespans, however, the underlying causes are still not completely understood^88^. Several predictive biomarkers of *C. elegans* aging have been described^89–92^ and the measurements of physiological processes, such as movement, pharyngeal pumping and reproduction have been used to predict lifespan^93^ and the age with an RMSE of 1.7 days^94^. A first transcriptomic clock of *C. elegans* aging using microarray data of 104 single wildtype worms predicted the chronological age with 71% accuracy^95^. When the prediction was based on modular genetic subnetworks inferred from microarray data with support vector regression, the age of sterile *fer-15* mutants at 4 timepoints was predicted with an r^2^ of 0.91. The same approach on the 104 individual N2 wildtype worms yielded an r^2^ of 0.77 indicating that for microarray data subnetworks of genes result in better prediction compared to single gene predictors, likely due to the noisiness of the datatype^96^. Although the accuracy of this model is reasonable, it is limited by the fact, that no lifespan-affecting genotypes or treatments were tested and that the validation dataset, although tested on single worms, resulted in an increased prediction error. Recently, an initial age prediction based on microarray data predicted 60 RNA-seq samples with a Pearson correlation of 0.54 and was improved to an r of 0.86 when the chronological age was rescaled by the median lifespan of the corresponding sample^97^. Even though this model instead of chronological age predicted the biological age of a variety of *C. elegans* genotypes, it is limited by the accuracy of the prediction. Moreover, the biological age is not reported in days, but as a variable with values between 0 and ∼2.5, which makes it harder to interpret.

To date, no aging clock for *C. elegans* has been built solely on RNA-seq data and been shown to predict the biological age of diverse strains, treatments and conditions to a high accuracy. In this study, we build such a transcriptomic aging clock that predicts the biological age of *C. elegans* based on high throughput gene expression data to an unprecedented accuracy. We combine a temporal rescaling approach, to make samples of diverse lifespans comparable, with a novel binarization approach, which overcomes current limitations in the prediction of the biological age. Moreover, we show that the model accurately predicts the effects of several lifespan-affecting factors like insulin-like signaling, a dysregulated miRNA regulation, the effect of an epigenetic mark, translational efficiency, dietary restriction, heat stress, pathogen exposure, the diet and dosage dependent effects of drugs. This combination of rescaling and binarization of gene expression data therefore allows for the first time to build an accurate aging clock that predicts the biological age regardless of the genotype or treatment. Lastly, we show how our model has the potential to improve the prediction of the transcriptomic age of humans and might therefore be universally applicable to assess biological age.

## Results

### Temporal scaling and transcriptome data binarization allow precise biological clock predictions

We downloaded and processed 904 publicly available RNA-seq samples for adult *C. elegans* for which corresponding lifespan data was available (Fig. 1, Table S1). Out of the 904 samples most (413) were wildtype N2 worm populations. A significant portion of 171 samples contained reads of temperature-sensitive sterile strains like *glp-1* or *fem-1* or double mutants thereof. 59 samples contained a mutation in the insulin-like growth factor 1 receptor *daf-2* and 45 a mutation in the dietary-restriction mimic strain *eat-2* either as a single or as a combination with a different mutation. 216 samples did not cluster in one of the mentioned groups and contain a variety of different strains. 112 of the samples span 14 different RNAi’s in 51 samples and 61 empty vector controls. Slightly more than half of the samples (490) were sequenced from a population that was undergoing a treatment (excluding RNAi or empty vector) that is different from the standard treatment of an *E. coli* OP50 diet at 20°C. The convoluted circle plot on the left side of Fig. 1 shows the overlap of the different possible combinations of strains, RNAi and treatments in our samples.

**Figure 1.**
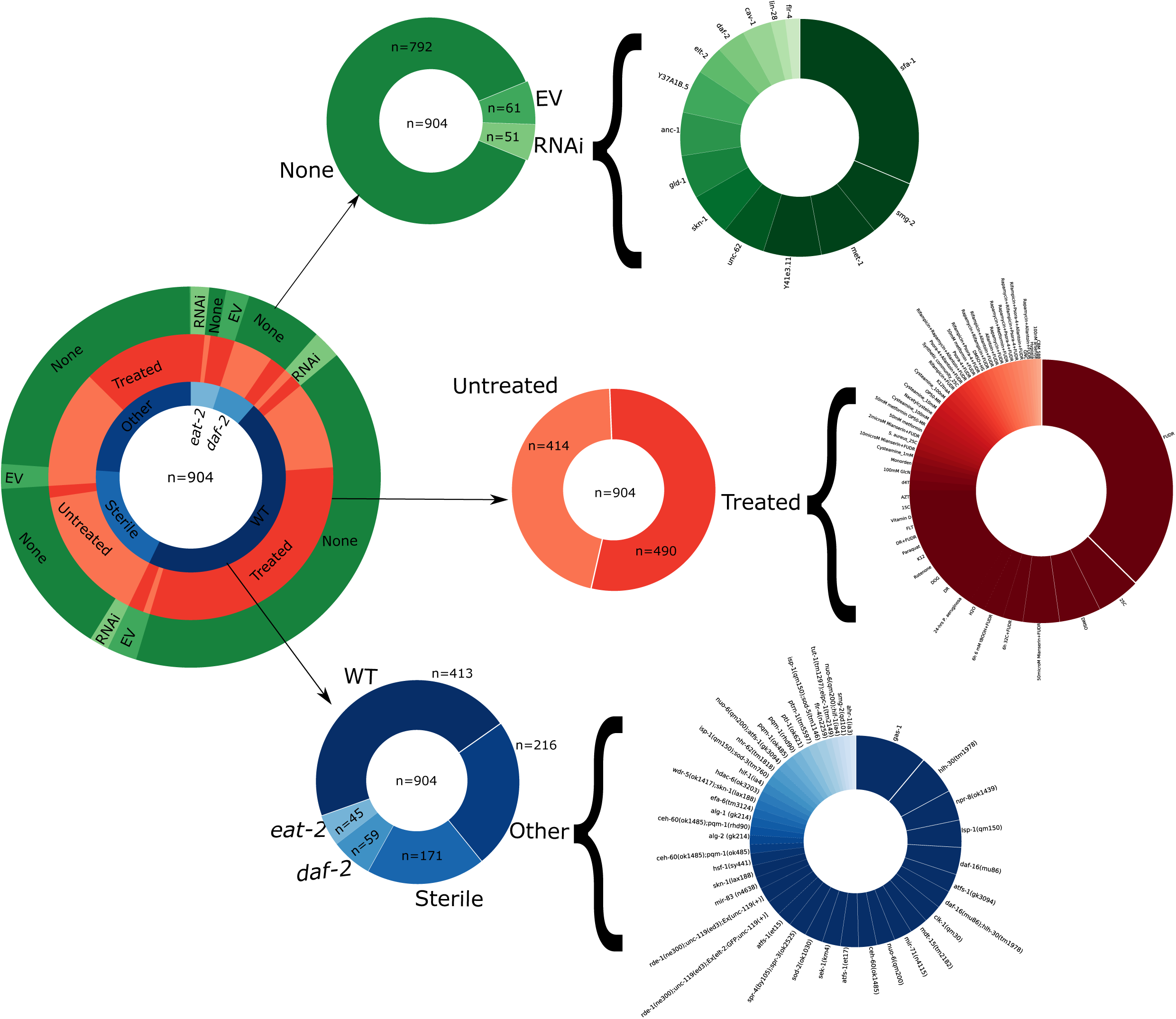
Data Overview. Overview of the processed published data utilized in the paper. Pie charts show the distribution of different genotypes (blue), treatments (red) and RNAi’s (green). The convoluted pie chart on the left shows the overlap of the three classes. The middle pie charts show broader clusters with the number of samples annotated. The partition ‘Sterile’ contains multiple different genotypes that can’t give rise to progeny and *daf-2*, as well as *eat-2*, might contain additional mutations. On the right finer partitions are shown for the RNAi’s in green, treatments in red and strains excluding WT, *eat-2, daf-2* and sterile mutants in blue. For a more detailed view see the Table S1.

We only downloaded and processed data for which the corresponding publication reported a median lifespan. The lifespan data is required to make strains with vastly different lifespans comparable. Without rescaling, an RNA-seq sample of a long-lived nematode beyond the normal lifespan of a wildtype worm would not be comparable to a wildtype sample, since no sample would be able to be generated. Lifespan-altering manipulations, e.g. a temperature shift, a *daf-2* mutation or oxidative damage, were shown to just shift the lifespan curve by stretching or shrinking it^98^. One interpretation would be, that all lifespan-affecting interventions converge on similar pathways, which affect the risk of death in a similar pattern, just at different velocities. Moreover, there have been descriptions of a transcriptional drift during *C. elegans* aging^97,99^, which might be due to a (dys-)regulation of single transcription factors^100^ and the suppression of this transcriptional drift might slow down the aging process^50^. Notably, age prediction could be improved by rescaling the chronological age by the median lifespan^97^.

We, therefore, employed a strategy similar to Tarkhov et al. and rescaled the lifespan by the corresponding median lifespan of the sample. We set the median lifespan of a standard wildtype N2 worm to µ=15.5 days adulthood. Using this standard lifespan, we calculated a correction factor to determine the biological age of a sample. For example, the correction factor of a strain with a measured median lifespan of 31 days, would be µ/31 = 0.5 and thereby assuming an aging rate reduction of 50%. This correction factor would be applied to each RNA-seq sample of the same strain and experiment. A sample sequenced e.g. at day 10 of adulthood, would be corrected to 10*0.5 = 5 days of biological age. Applying the individual correction factors for each RNA-seq sample, allows us to build a classifier of the biological, instead of the chronological age. Importantly, by defining a standard lifespan of 15.5 days we are able to predict the biological age in days instead of a variable between 0 and 2.5 as reported by Tarkhov et al.

Owing to the fact that the public data was generated in multiple laboratories with different protocols and sequencers (see Table S1 for details) we expected noisy data with a strong batch effect. Indeed, the results of an elastic net regression (see Methods for details) on the raw counts-per-million (CPM) reads, resulted in a mediocre model with an r^2^ of 0.78 a mean absolute error (MAE) of 1.02 days and a median absolute deviation (MAD) of 0.71 days. Fig. S1 shows the comparison of the rescaled biological age of the strains on the x-axis and the age predicted by the elastic net regression on the y-axis. Interestingly, the overall absolute error and the variance in the absolute error of the prediction increases strongly after ∼5 d (Fig. S2).

In order to mitigate this increase in variance, we used a novel approach and binarized the transcriptome data by setting the value of each gene to 1, if the CPM is bigger than the median CPM of the corresponding sample and 0 otherwise (see Methods for details), thereby reducing the noise, but retaining the information whether a gene is strongly transcribed or not. After this binarization, we trained an elastic net regression model with nested cross validation to obtain the best parameter setting and optimal set of genes (see Methods for details) that predict the biological age remarkably well with an r^2^ of 0.96, a MAE of 0.46 d and a MAD of 0.33 d (Fig. 2a).

**Figure 2.**
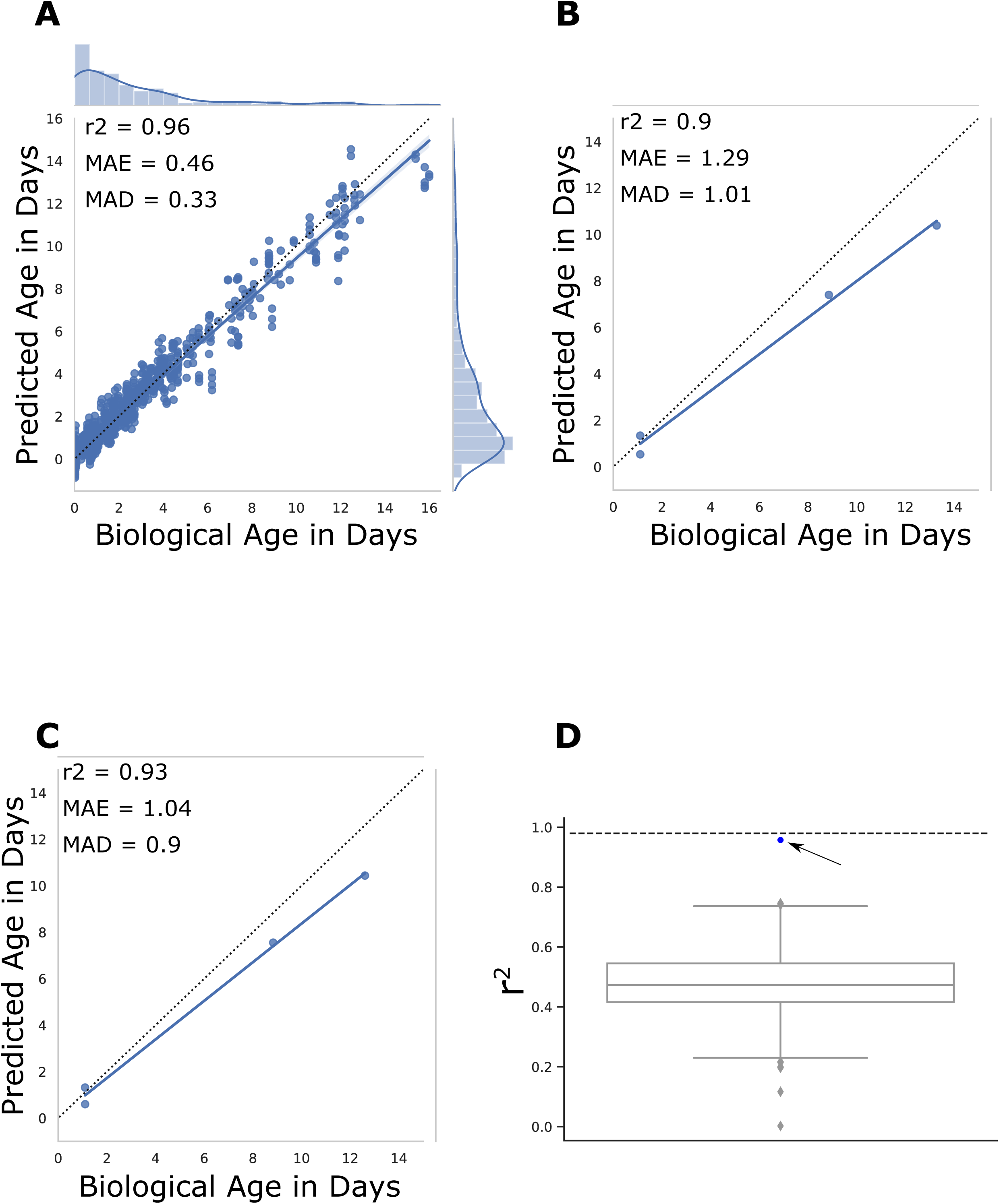
Biological age prediction. (A) Results of the biological age prediction computed by cross-validation. The x-axis shows the rescaled biological age in days starting from adulthood. The y-axis shows the predicted age computed by an elastic net regression on binarized gene expression data. Every blue dot displays one RNA-seq sample. The regression line is shown in blue and the dotted line shows the perfect linear correlation. The distribution of the data is shown on the side of the plot. MAE= mean absolute error, MAD= median absolute deviation. (B) Prediction of the model on a new independent dataset consisting of 4 WT samples at different time points. The x-axis shows the rescaled biological age in days starting from adulthood. The y-axis shows the predicted age computed by an elastic net regression on binarized gene expression data. Data from GSE65765^169^. (C) Prediction of the same set of samples improved by the second rescaling approach. (D) The y-axis shows the r^2^ of a given prediction. The box plot displays 1000 random models with 576 genes. The prediction by our final model with an r^2^ of 0.96 is shown as a blue dot and indicated by the arrow. The dotted line shows the theoretical limit of prediction given by the limit of accuracy in the chronological age annotation as well as variance in the lifespan data used for rescaling.

Interestingly, especially the increased variance in older samples, as seen in our initial analysis in Fig. S1, diminished and showed a strong improvement in overall accuracy. Comparison of the absolute error terms of the raw CPM and the binarized data prediction shows that the absolute error of the binarized prediction is lower than the prediction based on the raw CPMs regardless of the biological age of the worms. Furthermore, while the initial prediction on the raw data starts to get especially inaccurate starting from day 5, the increase in the binarized data is far less pronounced (Fig. S2a). Interestingly, also the variance of the absolute error terms stays more stable in the binarized data than the raw data and thereby demonstrating a more robust prediction regardless of the true age of the worms (Fig. S2b).

These results show that the binarization approach strongly improves the prediction, especially in older samples, which have been shown to contain a noisier transcriptome. Indeed, this age-dependent noisiness so far hindered the identification of proper aging biomarkers. The binarization therefore might facilitate the identification by reducing the noise, while retaining the important information. To verify our prediction further, an independent dataset, not used in the nested cross-validation for optimization of the parameter and gene set, was predicted with an r^2^ of 0.9, a mean error of 1.29 d and a median error of 1.01 d (Fig. 2b).

The results show that the overall prediction is highly accurate, however, although lower than the increase in deviation in the raw data, the binarized data as well show a decrease in accuracy in samples with an older biological age (see also Fig. S2). This might be due to the lower sample size of older animals, but might also be influenced by the nature of bulk RNA-sequencing itself. Fig. S3a shows a standard lifespan curve of *C. elegans*. Until ∼day 8 100% of non-censored worms are alive. Starting from day 8 the first worms die, until the median lifespan is reached at ∼15.5 days and the maximum at ∼24 days. We can assume that the biological age of worms at the same chronological age follows a normal distribution (Fig. S3b). In other words, in a plate of synchronized worms at day 8 we would expect to see that most worms are also at a biological age of 8 days. However, some worms will be healthier while others are already close to death and will therefore be the worms that start dying early. While the peak of this bell curve will therefore be the chronological age of the worm population, some worms will be biologically younger and some older (Fig. S3b). Starting from the next day, the first part of the worm population will die (Fig. S3c). Assuming the normal distribution of the biological age of the worms and a hypothetical maximum biological age as shown with the dotted line in Fig. S3d we can hypothesize that the biologically older worms will die off first and thereby truncate the biological age distribution on the right side of the curve (Fig. S3d). This truncation will shift the true median biological age towards the left side, as indicated by the green line. This becomes more noticeable at the median lifespan of 15.5 days, where by definition 50% of the population is dead (Fig. S3e). Following the same reasoning from above, we see that the right half of the biologically older worms died, while the younger half of the population stayed alive. However, this clearly skews the distribution, since the oldest 50% of the population is dead and therefore won’t contribute to the average biological age anymore. Indeed, the median biological age, will be the median of the remaining, alive worms, i.e. the left part of the curve. This will result in a shift of biological age, especially for chronologically older populations (Fig. S3f). In consideration of this biological age shift, an RNA-seq sample sequenced at 15.5 days will have a younger true population-median biological age, which will introduce a bias into the regression model. The bias will be not as pronounced in younger samples, since most of the population will still be alive (Fig. S3b).

To alleviate this bias, we calculated a second correction term that takes into consideration the hypothetical biological age distribution of the sequenced population (methods for details). Applying this correction before the optimization of the regression, resulted in an improved prediction model, especially for the independent dataset. The new model utilizes 576 genes and predicts the full dataset slightly better, with an r^2^ of 0.96, a mean error of 0.45 d (−1.73 %) and a median error of 0.32 d (−2.18 %) (Fig. S4a). The independent dataset is now predicted with an r^2^ of 0.93, a mean error of 1.04 d (−19.55 %) and a median error of 0.9 d (−11.57 %) (Fig. 2c). These data indicate that it might be worthwhile including a correction for the survival bias of worms in older populations.

To confirm that not every gene set of 576 genes results in a similar prediction, we randomly sampled 576 genes and recorded the resulting absolute errors and r^2^ values. The boxplot in Fig. 2d shows the distribution of r^2^ values centering around the mean of 0.488 with a standard deviation of 0.117. The blue dot shows the result of our predicted gene set as a clear outlier at 0.96. The MAE and MAD are centered around 1.27 d and 0.911 d with a standard deviation of 0.066 and 0.063 respectively (Fig. S4b).

To assess the precision of the age prediction, we next probed how close this model approaches the theoretical limit of a biological clock. The downloaded data is annotated in whole days alive from adulthood and thereby including a variance of +/-12 h to the actual chronological age. Random sampling of this error alone gives a mean error of 0.236 (+/-0.006) d, a median error of 0.187 (+/-0.006) d and a r^2^ of 0.986 (+/-0.002). However, since lifespan assays, even done under the same conditions in the same laboratory, will vary, we can assume that the reported median lifespan, used for the temporal rescaling, will also be including an inherent experimental error. Indeed, it has been shown that lifespan assays are heavily affected by the number of animals and less, but substantially, by the scoring frequency, thereby indicating that many lifespan studies are underpowered and often driven by stochastic variation^101^. Computing the mean and SD of lifespan assays for one genotype with the same treatment for several publications, shows that the variation is indeed on average ∼7 % for one standard deviation from the mean with a range between 5.44 % to 8.83 % (Table S3). An assumption of a moderate 5 % deviation between assays increases the mean error to 0.302 (+/-0.007) d, the median error to 0.244 (+/-0.008) d and reduces the r^2^ to 0.98 (+/-0.002). These theoretical optima, shown as dotted lines in the boxplots in Fig. 2d and Fig. S4b, clearly display the quality of our prediction. We conclude that the prediction based on the set of 576 genes is close to the theoretical optimum.

Next, we compared our model to a previous model^97^ that described three sets of aging-associated genes. The first set, consisting of 327 genes was generated by a meta-analysis of publicly available microarray data, the second consists of 902 age-associated genes generated by an RNA-seq experiment, and finally, a sparse subset with only 71 genes that was used for the biological age prediction. The gene set derived from microarray data performed worst on the prediction of the 904 RNA-seq samples with an r^2^ of 0.52, followed by the gene set of 902 genes with an r^2^ of 0.58 and finally the sparse predictor with an r^2^ of 0.62 (Fig. S5a-c). The latter corresponds to a Pearson correlation of 0.79, and is thereby similar to the Pearson correlation of 0.86 as published by Tarkhov et al. Binarization improves the prediction of the two larger gene sets as well to an r^2^ of 0.74 and 0.78 respectively (Fig. S5d-e). The r^2^ of the sparse predictor decreased to 0.57, however, the MAE and MAD decreased and thereby also show that a single quality assessment is not enough to give a good evaluation (Fig. S5f).

These comparisons indicate that our new model consisting of 576 binarized genes predicts the biological age of *C. elegans* to a high accuracy and superior to previously existing models.

### Prediction of multiple lifespan-affecting factors

Since our model is able to predict the biological age to a high accuracy, we next tested the capability of the model to predict the effect of multiple lifespan-affecting factors. We used the previously determined 576 predictor-genes and trained an elastic net regression on the 904 RNA-seq samples, excluding the data for the respective publication. This is thereby a different cross-validation approach where we excluded a whole dataset from a paper at a time.

First, we tested the well-known effect of insulin-like signaling on the biological age and could see that a *daf-2* mutation reduces the predicted biological age compared to the WT strain of the same experiment significantly by 41.3 % in 4-day adult *C. elegans* (Fig. 3a). This corresponds to a 1.71-fold lifespan extension. Since the WT sample of this dataset^102^ was already longer lived than our standard 15.5 days, we computed also the comparison against 15.5 d which resulted in a 2.31-fold increase in lifespan for *daf-2*. The even longer lived *daf-2; rsks-1* double mutant is accordingly predicted to be even younger with a significant reduction of 56.8 % in 4-day adults, corresponding to a 2.32-fold lifespan extension (Fig. 3b)^103^.

**Figure 3.**
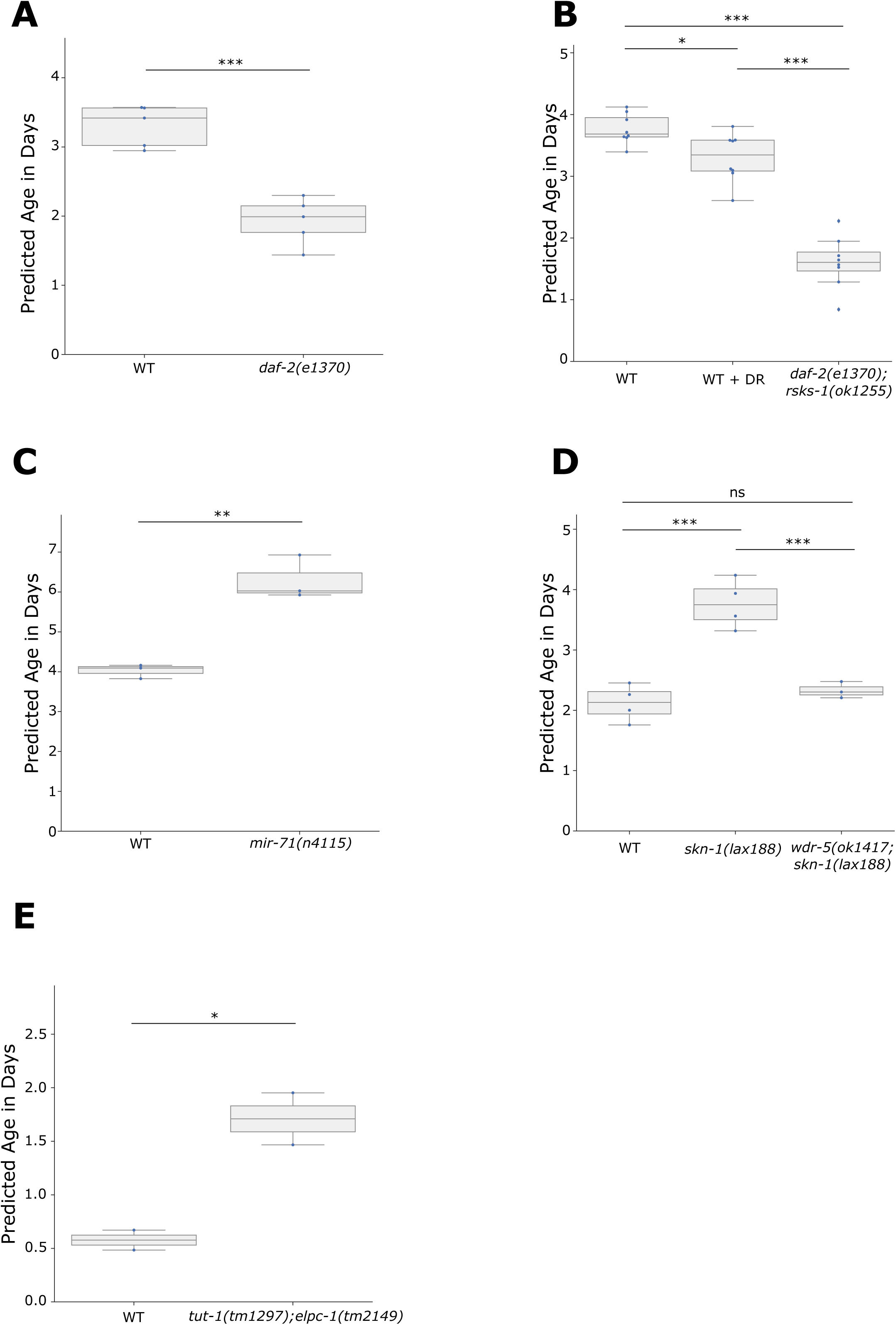
Biological age prediction of short- and long-lived mutants. The box plots show the predicted biological age in days on the y-axis. Assuming the properties of a uniform temporal rescaling a lower predicted age will equal a longer lifespan. The corresponding whole dataset was set aside for the training of the final model for the corresponding plot. Blue dots display single RNA-seq samples. (A) The lifespan-extending *daf-2(e1370)* strain is predicted to be biologically younger than WT samples of the same chronological age (4.5 days). Note that the WT strain in this publication had a longer lifespan (19.4 days) than the standard 15.5 days and is thereby also predicted to be biologically younger than its chronological age. Data from GSE36041 ^102^. (B) Dietary restriction (DR) and the long-lived double mutant *daf-2(e1370); rsks-1(ok1255)* are predicted to be significantly younger than WT samples of the same chronological age (4 days). Data from GSE119485 ^103,170^. (C) The lifespan-shortening *mir-71(n4115)* mutation significantly increased the predicted biological age compared to samples of the same chronological age (5 days). Data from GSE72232 ^104^. (D) The gain-of-function mutant *skn-1(lax188)* significantly increased the biological age, while an additional mutation in the epigenetic regulator *wdr-5* rescues the biological age back to WT levels (2 days). Data from GSE123531 ^105^. (E) The double mutant *tut-1(tm1297); elpc-1(tm2149)* significantly increases the biological age (chronological age of 1 day). Data from GSE67387 ^106^. *p<0.05, **p<=0.01, ***p<=0.001, independent t-tests were used for comparisons in (A), (C) and (E). One-way ANOVA with a post-hoc Tukey test was used in (B) and (D). Table S4 contains more detailed statistics.

To see, whether short-lived mutants can also be predicted correctly, we next tested *mir-71*, which has been shown to regulate the global miRNA abundance during aging and to directly influence lifespan^104^. Compared to wildtype, *mir-71* is predicted to be 56 % older in 5-day adults, corresponding to a -1.56-fold lifespan reduction (Fig. 3c). In addition, samples of a gain-of-function *skn-1* mutation, detrimental for lifespan, got correctly predicted to be 77.2 % older than wildtype worms at day 2 (Fig. 3d). Interestingly, this adverse effect can be rescued by a loss-of-function mutation in *wdr-5* and the subsequent abolishment of the epigenetic mark H3K4me3^105^, which is remarkably also reflected in our prediction. Loss of protein homeostasis decreases overall fitness and is a hallmark of aging. In *C. elegans* the loss of uridine U34 2-thiolation in *tut-1; elpc-1* double mutants has been shown to have a negative impact on efficient translation and to promote protein aggregation^106^. Strikingly, this effect on translational efficiency is also reflected in the transcriptomic aging clock for day-1 adults, which are predicted to be 196 % older than their wildtype counterpart (Fig. 3e).

These data show, that the transcriptomic clock can effectively predict the biological age of a variety of mutants and pathways, ranging from the insulin pathway, miRNA’s, the epigenetic mark H3K4me3 and translational efficiency.

Since both, long-lived and short-lived strains seem to be predicted with the correct pattern, we next asked whether we could predict the effect of dietary restriction (DR) on the biological age. Although a slight effect, the dietary restricted worms are predicted to be 12.9 % younger than their normal-fed counterpart at day 4 of adulthood (Fig. 3b). Dietary restriction induced longevity was shown to depend on the PMK-1/p38 signaling-regulated innate immune response. In *C. elegans sek-1* is part of the PMK-1/p38 signaling cascade and required for longevity in dietary restricted worms^107^. Noticeably, the same trend can be observed in our prediction for day-6 adults (Fig. 4a). A two-way ANOVA showed a significant interaction between the effects of the strain and dietary restriction (p=0.004), which indicates that the effect of DR is dependent on *sek-1* activation. Although in this dataset the adjusted p-value of the effect of DR in WT worms is not significant (p=0.057) it is interesting to note, that the dietary restricted worms are on average 32 % younger than the ad libitum fed WT worms and thereby showing a stronger effect than the 12.9 % reduction in Fig. 3b. This could be due to strain differences of the different laboratories or suggest that positive effects of DR add up over time.

**Figure 4.**
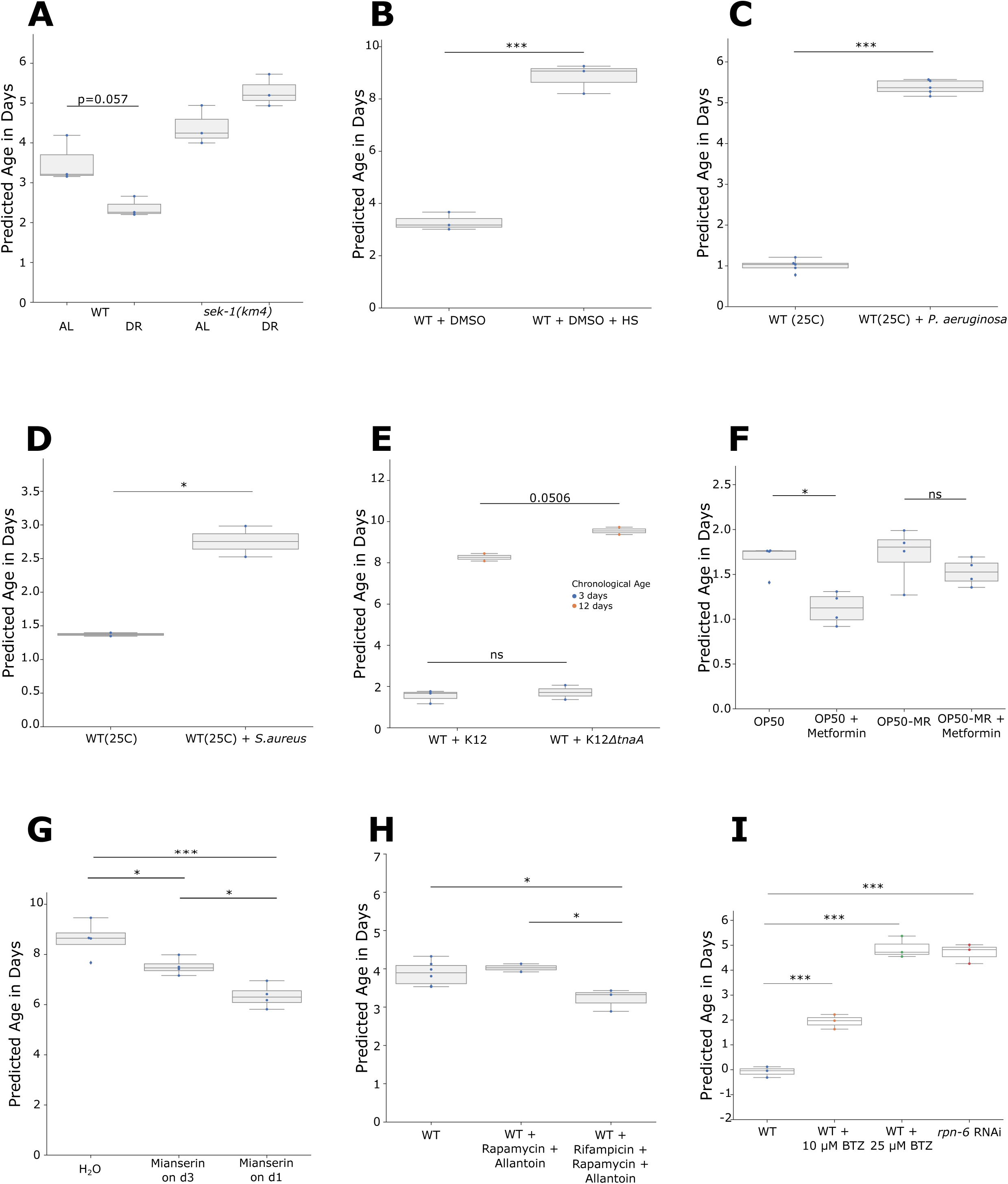
Biological age prediction of a variety of treatments and stressors. The box plots show the predicted biological age in days on the y-axis. Assuming the properties of a uniform temporal rescaling a lower predicted age will equal a longer lifespan. The corresponding whole dataset was set aside for the training of the final model for the corresponding plot. Blue dots display single RNA-seq samples. (A) The genotype-dependent effect of dietary restriction (DR) is resembled in the prediction of chronologically 6-day adults. A two-way ANOVA shows a significant interaction effect (p=0.004) between the genotype and the diet. AL = ad libitum fed. Data from GSE92909 ^107^. (B) Heat shock induces a strong increase of the predicted biological age at a chronological age of 3 days in WT. Data from PRJNA523315 ^55^. (C) Pathogen infection by *P. aeruginosa* at 25 °C at chronological age day 1 increases significantly the predicted age. Data from GSE122544 ^171^. D) Pathogen infection by *S. aureus* at 25 °C at chronological age day 1 increases significantly the predicted age. Data from GSE57739 ^172^. (E) The change in diet from K12 to K12*ΔtnaA E. coli* shows an increasing trend, especially in chronologically older population, as indicated by the different colors. A two-way ANOVA shows a significant diet effect (p=0.03) and almost significant interaction effect (p=0.067). Data from GSE101910 ^108^. (F) The bacterial-strain-dependent effect of Metformin is resembled in the prediction. The box plots show wildtype worm populations at a chronological age of day 2 with either a standard OP50 *E. coli* diet or a Metformin-resistant OP50 (OP50-MR) strain with or without 50mM Metformin. A two-way ANOVA showed a significant treatment effect (p=0.004). Data from E-MTAB-7272 ^109^. (G) The dosage-dependent effect of Mianserin is resembled in the prediction. The box plots show wildtype worm populations at a chronological age of day 10 either treated with water or 50µM Mianserin on day 3 or day 1. A one-way ANOVA showed significance (p=0.0008). Data from GSE63528^50^. (H) The effect of drug combinations at the chronological age of 6-days is resembled in the prediction. A one-way ANOVA showed significance (p=0.02). Data from GSE108263 ^110^. (I) An independent dataset without a reported lifespan sequenced at the chronological age of day 1. Wildtype worms were treated with either 10 µM or 20 µM of the proteasome inhibitor Bortezomib (BTZ), or RNAi against the proteasomal subunit *rpn-6*. Data from GSE124178 ^111^. *p<0.05, **p<=0.01, ***p<=0.001, independent t-tests were used for comparisons in (B), (C), and (D). One-way ANOVA with a post-hoc Tukey test was used in (G), (H), and (I). Two-way ANOVA with a post-hoc Tukey test was used in (A), (E), and (F). Table S4 contains more detailed statistics.

Next, we decided to test, whether different lifespan-shortening stressors can be predicted correctly. Both heat stress (Fig. 4b) and pathogen exposure to either *P. aeruginosa* or *S. aureus* (Fig. 4c, d) showed a strong increase in the predicted biological age. The former increased the prediction by 169.3 % in day 3 adults, while the latter increased the predicted age by 421.4 %, respective 101 %, in day 1 adults. While heat or pathogen exposure can lead to a quick demise of the animals, we wondered whether also more subtle changes in lifespan by different diets and subsequent nutrient metabolism could be detected. It has been shown that an *E. coli* K12 variant’s indole secretion extends fecundity and overall health- and lifespan in *C. elegans*, while an isogenic *E. coli* strain (K12tnaA) with a deletion in the indole-converting gene does not have these benefits. This effect on health span was reported to be not yet visible in worms on day 8, but only showed a significant difference on the next tested timepoint on day 15 ^108^. Intriguingly, the same pattern can be observed in RNA-seq samples of day 3 and day 12 (Fig. 4e). A two-way ANOVA showed a significant treatment effect (p=0.034) indicating the sensitivity of the approach. Moreover, in accordance with the published results, a subsequent Tukey post-hoc test showed no difference between the diets on day 3 (adjusted p=0.9), while day 12 showed a 15.3 % increased biological age in the K12tnaA diet (adjusted p=0.0506). Consistent with the link between diet-dependent changes in nutrient metabolism and lifespan, it has been shown that the lifespan-extending effect of Metformin is, at least partially, regulated by a bacterial nutrient pathway^109^. A two-way ANOVA of the predicted biological age of day-2 adults, grown on either *E. coli* OP50 or a Metformin-resistant OP50 strain, with or without Metformin showed as well a significant bacteria effect (p=0.045) as a significant drug effect (p=0.004). A subsequent Tukey post-hoc test showed a significant reduction of the biological age of Metformin treated wildtype worms grown on OP50 (−34.5 %), but no significant effect in worms grown on Metformin-resistant OP50 (Fig. 4f).

Next, we asked whether the effect of the duration time of a drug, might be reflected on the transcriptomic age. The antidepressant Mianserin has been shown to extend the lifespan of *C. elegans* by inhibiting serotonergic signals, which is lessening the age-dependent transcriptional drift. This effect is more pronounced in animals that were treated starting from day 1, compared to starting the treatment from day 3 ^50^. Our prediction of day 10 adults resembles this conclusion; a one-way ANOVA showed a significant difference (p=0.0008) and an ensuing Tukey post-hoc test revealed statistical significance between all three cases, with the biggest effect in worms treated from day 1 (Fig. 4g).

An interesting and challenging questions is, whether the combination of different lifespan-extending drugs might have a synergistic effect or not. Admasu et al. reported that not all combinations of drugs have an additional effect. While the combination of Rapamycin with Allantoin had no effect on the lifespan of wildtype worms, the triple combination with Rifampicin surprisingly had the biggest effect^110^. Interestingly, while the administration of Rifampicin, Rapamycin and Allantoin significantly reduced the predicted age by 17.7 % (Fig. 4h), the double combination of Rapamycin and Allantoin did not change the predicted lifespan, which is in accordance with the published lifespan results.

Lastly, we decided to check the effect of proteotoxic stress on the transcriptional age and downloaded a new dataset for which no direct lifespan data was published and which contained treatments that were not included in any of the analyses and nested cross-validations above^111^. We tested the samples of two different dosages of the proteasome inhibitor bortezomib (BTZ) and the knockdown by RNAi of the proteasomal subunit RPN-6.1 and saw a significant increase in the biological age or all three samples (Fig. 4i). Notably, the effect of BTZ shows a dose dependency. *rpn-6*.*1* RNAi has been shown to strongly reduce the lifespan of WT worms^112^ and BTZ supposedly mimics the effects by directly blocking the proteasome and has been shown to dramatically reduce the lifespan of starved worms^113^. Moreover, although no direct lifespan data is available for normal fed worms, 10 µM BTZ leads to an early death starting from day 3 ^111^, while 25 µM even increased the dying rate (Fabian Finger, personal communication). These results demonstrate that the nested cross-validation was sufficient to prevent overfitting, that our model extends beyond the data described here and that even lifespan-affecting stressors unknown to the model, i.e. proteasomal stress, are correctly predicted.

In conclusion, we demonstrated that our transcriptomic aging clock of *C. elegans* is highly accurate and versatile usable. We showed, that it correctly predicts the effects of insulin-like signaling, a modified miRNA regulation, the effect of an aberrant active transcription factor and the reversal of this effect by an epigenetic mark, translational efficiency, dietary restriction and the requirement of the intact innate immune system on its lifespan-extending effect, heat stress as well as pathogen exposure and the effects of diet-depending metabolites. Lastly, we also showed that the predictor is able to correctly identify the effect of Metformin through the hosts microbiota, the dosage-dependent effect of drugs and the counterintuitive fact that the combination of lifespan extending drugs might not be necessarily synergistic. Strikingly, our model extends beyond the data used for the nested cross-validation and is able to correctly predict the biological age of worms, for which no direct lifespan data was available.

### Functional characterization

The final regression model utilizes 576 genes, out of which 294 have a negative coefficient and thereby are mostly expressed in young worms, while 282 genes have a positive coefficient and thereby increase the predicted age if active. Intriguingly, the protein-coding genes with a negative coefficient were enriched on the X-chromosome and are significantly less expressed from chromosome I and II (Fig. S6a). Protein-coding genes with a positive coefficient show a opposite trend and are significantly enriched on chromosome I and II, while depleted from chromosome IV (Fig. S6b, c). Interestingly, a WormExp^114^ gene set enrichment analysis of the genes with a negative coefficient, so those that are associated with younger samples, are enriched in age-related categories that are downregulated with aging (Fig. 5a). Moreover, the 294 genes are enriched in the *pmk-1, elt-2, pqm-1* and *daf-16* transcription factor target category (Fig. 5b). A motif search at the promoter regions of the genes with a negative coefficient corroborates this finding and shows a significant enrichment in the GATA transcription factors PQM-1 and ELT-3 (Fig. S7a). Although the gene set enrichment analysis with WormExp did not show a significant enrichment of transcription factors in the gene set with a positive coefficient, the motif search also identified the GATA motif enriched at the promoter regions (Fig. S7b). Notably, the GATA transcription factor *elt-6* is within the top 30 % of genes with a positive coefficient in our gene set and thereby correlated with older worms and has been shown to increase during normal aging and to increase the lifespan upon knock down by RNAi^115^. Interestingly, genes associated with younger worms are also enriched in genes that are upregulated in germline-ablated animals (Fig. 5c), which in general exhibit an increased lifespan. Genes with a positive coefficient on the other hand are enriched in categories that show an increase with age (Fig. 5d).

**Figure 5.**
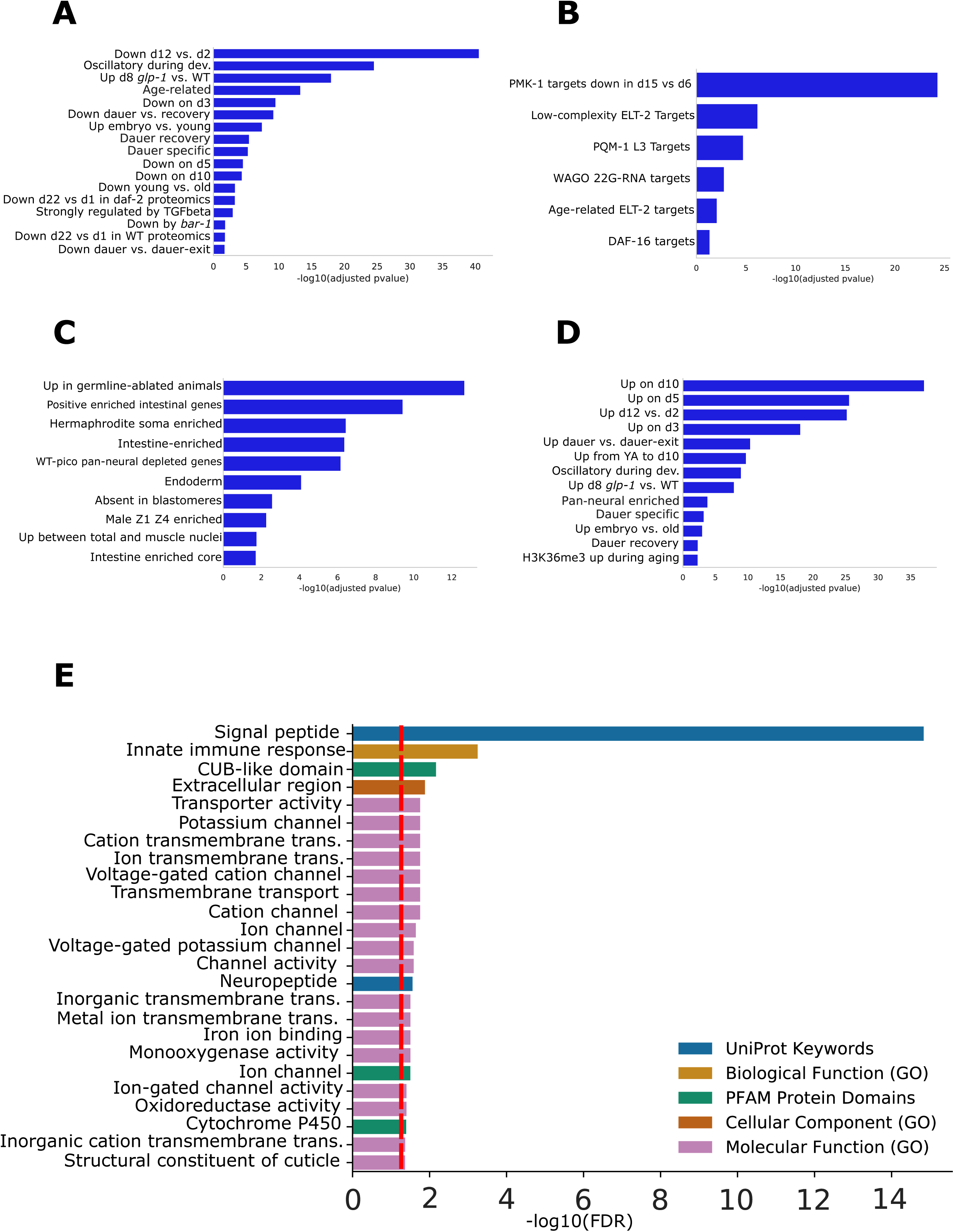
Functional analysis of the predictor genes. (A-D) WormExp gene set enrichment analysis for the 576 predictor genes. The x-axis displays the -log10 of the adjusted p-value. Only statistically significant (adjusted p<0.05) enrichments are shown. (A-C) Gene set enrichment analyses for the genes with a coefficient <=0 for the Development/Dauer/Aging category (A), the TF Targets category (B), and the Tissue category (C). (D) Gene set enrichment analyses for the genes with a coefficient >0 for the Development/Dauer/Aging category.nt analyses for the gene(F) Functional enrichment analysis for the 576 predictor genes by String and geneSCF. The x-axis displays the -log10 of the adjusted p-value. The red line displays an adjusted p-value of 0.05. Different enrichment categories are color-coded.

A subsequent functional enrichment analysis with String v.11^116^ and geneSCF^117^ revealed a strong enrichment of signal peptides, transporter activity, and neuropeptides, which suggest that especially systemic responses influence the aging process (Fig. 5e). Neurotransmitters, although not directly enriched in the GO-term analysis, might as well play an important role. *hic-1* is one of the genes with the strongest increase in predicted age of our gene set. It has been previously shown to be present at the presynaptic terminal of cholinergic neurons and to regulate the normal secretion of acetylcholine neurotransmitter and Wnt vesicles^118^. In the same manner, the dopamine receptor *dop-4* is in the top 25 % of genes with a negative coefficient and has been shown to promote healthy proteostasis and the innate immunity as well as detoxification genes^119^. Interestingly, the innate immune response and cytochrome P450 enrichment in our gene set might indicate a role of a general stress response, detoxification and drug metabolism during the aging process. Consistent with a general stress response, we also find *csa-1* in the list of genes with a positive coefficient, which might indicate an increased DNA damage load in older worms.

To conclude, these results further validate the genes used for the age prediction and indicate that the aging process might be driven by the dysregulation of single transcription factors (Fig. 5b) and a systemic signal transmitted by signal peptides (Fig. 5e).

### Human data

To demonstrate, that our novel approach is also usable for other organism we downloaded a recent human dermal fibroblast RNA-seq dataset generated from cell culture of 133 healthy individuals with ages between 1 and 94 and 10 patients with Hutchinson-Gilford progeria syndrome (HGPS) with ages between 2 and 9 ^61^. Fleischer et al. showed that an LDA ensemble approach can predict the age of the 133 healthy patients with a r^2^ of 0.81, a mean error of 7.7 years and a median error of 4.0 years. Moreover, they find a statistical increased predicted biological age of HGPS patients, as would be expected from a premature aging disease. However, as they mention, the ensemble method has some limitations, i.e. the discretization of age, the computational cost and the difficult interpretation of the influence of gene expression changes on the predicted age.

Our regression-based method is fast to compute, does not require the discretization of age and directly allows the effect-interpretation of the activity of single genes on the predicted age. Using the elastic net regression on the unbinarized data resulted in a model of 132 predictor genes and in a similar prediction quality as the elastic net regression by Fleischer et al. (Fig. S8a) and similarly the HGPS samples are not predicted to be biologically older (Fig. S8b). However, binarization of the data before calculating the elastic net regression improved the results dramatically to an r^2^ of 0.91, a MAE of 6.63 years and a MAD of 5.24 years (Fig. 6a). Moreover, our model predicts the HGPS patients to be significantly older (Fig. 6b). This new model contains 141 predictor genes (Table S5), out of which 25 are significantly enriched in the biological process regulation of cell death. Interestingly, among the predictor genes the forkhead transcription factor FOXO1 –a regulator of the aging process in *C. elegans* and mammals– is positively correlated with age thus further supporting the evolutionary conservation of transcriptionally regulated longevity mechanisms^120^.

**Figure 6.**
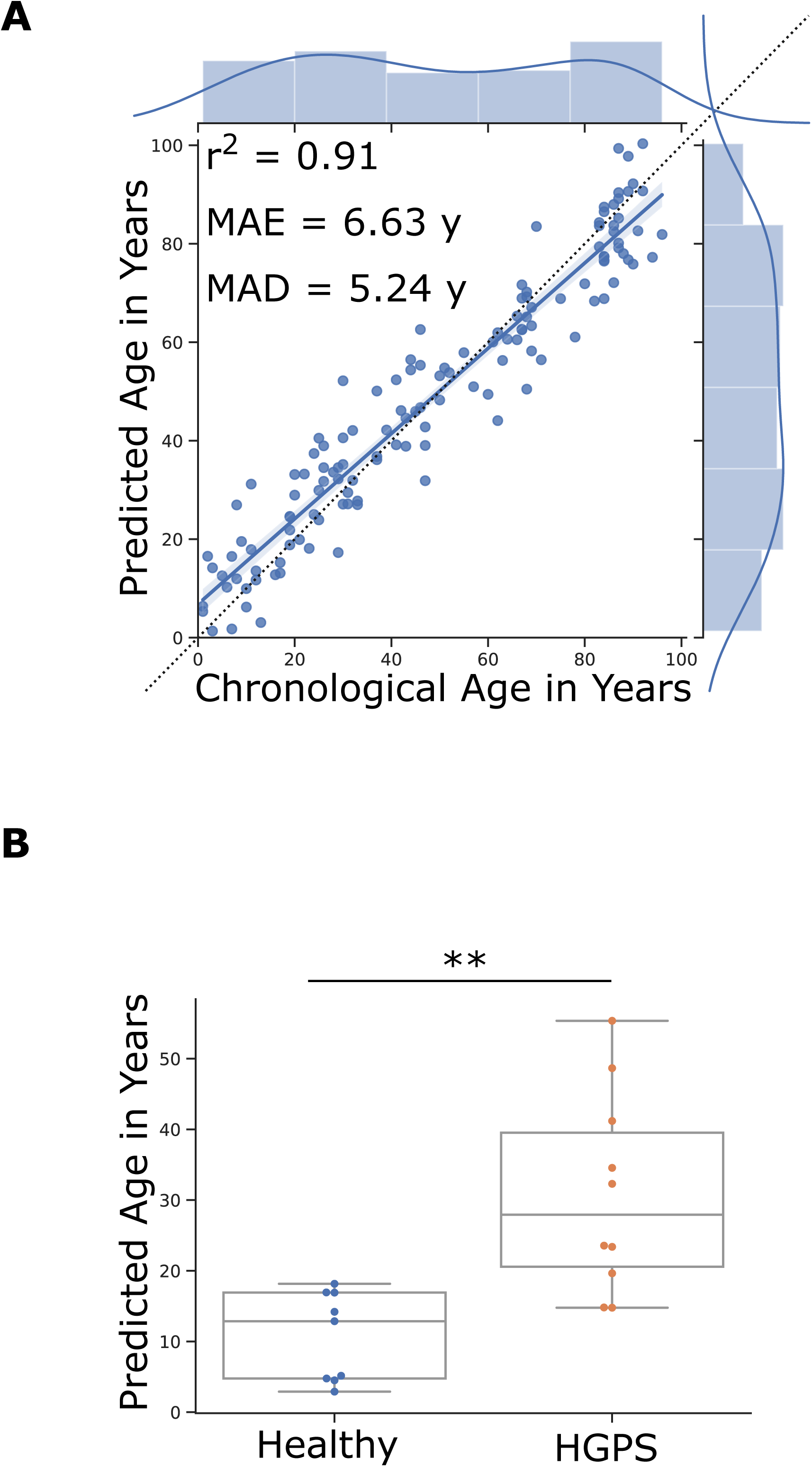
Transcriptomic human aging clock. (A) Results of the age prediction computed by cross-validation on human fibroblast gene expression data. The x-axis shows the chronological age in years. The y-axis shows the predicted age computed by an elastic net regression on binarized gene expression data. Every blue dot displays one RNA-seq sample. The regression line is shown in blue and the dotted line shows the perfect linear correlation. The distribution of the data is shown on the side of the plot. MAE= mean absolute error, MAD= median absolute deviation. Data from GSE113957 ^61^. (B) Box plots of age predictions of samples from Hutchinson–Gilford progeria syndrome patients (red) and predictions of age-matched healthy controls (blue) by the elastic net regression of binarized gene expression data. Progeria samples are predicted to be significantly older than age-matched healthy controls. **p<=0.01, calculated by an independent t-test. Table S4 contains more detailed statistics.

To summarize, these data indicate, that elastic net regression on binarized gene expression data is not only usable in the nematode *C. elegans*, but possibly also in more complex organisms like humans.

## Discussion

The molecular understanding of aging on the genetic^121^, epigenetic^122^, transcriptomic^123^, proteomic^69^ and metabolomic^124^ level has made steady progress over the recent years. Especially usage of *C. elegans* has led to a number of important discoveries^125,126^. However, up to date no single model could predict the biological age of any organism to a high accuracy in diverse strains, treatments and conditions. In our study, we show that the binarization of gene expression data allows a biological age prediction of *C. elegans* to an unprecedented accuracy and for the first time the prediction of a variety of lifespan-affecting factors. Additionally, we show that the binarization approach, even without the biological rescaling might be applicable to and improving the predictions of other organisms. This is in contrast to the currently most widely used epigenetic clocks, which are limited to organisms with DNA methylation marks. Moreover, our results suggest that especially the innate immune system and neuronal signaling are important for an accurate prediction and therefore also might play an essential role in the aging process.

Binarization of the gene expression data hugely improved the predictability of the biological age. Interestingly, the biggest deviation from the true biological age is in the samples treated with heat shock or in *mir-71, eat-2* and *skn-1(gof)* mutants. Heat shock treatment and an *eat-2* mutation have been shown to exhibit a different aging trajectory and to diverge from the temporal scaling approach proposed by Stroustrup^98^. Similarly, *skn-1(gof)* and *mir-71* display a sharp drop in lifespan^104,105^ that cannot totally be accounted for with our median-lifespan-rescaling approach. Incorporating the whole lifespan curve could therefore improve the prediction even further. In this regard, it is also noteworthy that the utilized bulk-sequencing data introduce several biases that might not be reflected in a simple rescaling approach. We tried to alleviate some of the potential biases with our second rescaling approach, which should reduce the error that is introduced by the fact that especially the biologically older part of a population dies of first. However, it has been published that *C. elegans* dies of at least 2 different types of death^127^. An early death with a swollen pharynx, induced by an increased bacterial content, or a later death with an atrophied pharynx. This might introduce a different bias, since the initial transcriptional response close to an early death might be different from the response to a later death. Nevertheless, even with these limitations our model predicts the biological age of worms remarkably well.

The increasing error and increase in variance of the age predictor in older worms is especially visible in the unbinarized model. This might be due to the known age-dependent increase in transcriptional variety^128,129^ that limits the ability of the regression model to pick an accurate subset of genes. Different hypotheses have been proposed that try to explain this transcriptional noise. In *C. elegans* it might be partially regulated by a microRNA feedback loop that is dependent on *mir-71*^104^, serotonergic signals^50^, and the decline of the GATA transcription factor ELT-2 during aging^100^. One interesting possibility is the idea that the increasing noise is driven by accumulating somatic mutations over the course of aging. Indeed, Enge et al. demonstrated an increase in the transcriptional noise as well as an age-dependent accumulation of somatic mutations in single human pancreatic cells, however, they did not find any support for a causal relationship between exonic mutations and transcriptional dysregulation^130,131^. Interestingly, it has been recently proposed that single-cell aging is split into a normal aging and a catastrophe aging phase. During the normal aging phase transcriptional noise even decreases, while it dramatically increases in the second aging phase^132^. The authors propose that chromatin state transition rates and thereby also the stability of the regulatory network might play an important role. Indeed, it has been shown that the stability of a gene network is intrinsically linked to longevity and genotoxic stress resistance^43^. An important factor of transcriptional regulation and therefore network stability are transcription factors.

### Transcription Factors

Similar to Tarkhov et al. we find an enrichment in targets of DAF-16, the GATA transcription factors PQM-1 and ELT-2, and PMK-1 in our predictor gene set. DAF-16 is known to be involved in a variety of stress responses and longevity pathways^133–136^. Furthermore, it has been shown to be activated during the normal aging process, where it regulates different targets from the canonical *daf-2*-mediated activation^137^. The authors proposed that this stabilizes the transcriptome during normal aging. GATA transcription factors have been found to be relevant for a variety of tissue-specific stress responses^138^, to have a functional role in the aging process^115^, and to promote together with *daf-16* developmental growth and survival amid persistent somatic DNA damage^139^. Moreover, deactivation of *elt-2* has been described as a major driver of normal *C. elegans* aging^100^ and *pqm-1* has been shown to decline with age, to be involved in *daf-2*-mediated longevity^140^ and reproductive aging^141^. Interestingly, it has been shown that *pqm-1* is required for the systemic stress signaling pathway upon proteotoxic stress via innate immunity-associated proteins^142^. The p38 MAPK family member *pmk-1* is an important gene in the nematode’s pathogen defense system and innate immunity.

### Innate Immune Response

The innate immune system of *C. elegans* has been linked to several lifespan-affecting pathways^143,144^ and a general systemic stress resistance^145^. Schmeisser et al. for example showed that dietary restriction (DR) dependent lifespan extension requires a limited neuronal ROS signaling via a reduced mitochondrial complex 1 activity that activates PMK-1/p38^146^. Conversely, a recent study showed, that DR extends lifespan dependent on the downregulated, but intact p38-ATF-7 pathway^107^. The lifespan extension from reduced insulin-like signaling has similarly been shown to be at least partially dependent on the p38 response to AMPK-induced ROS signaling^102^ and that this response works in parallel to the DAF-16 regulation of longevity^147^. Notably, the often-used DR-mimic *eat-2* mutants have also been shown to only display longevity if grown on a bacterial lawn that activates the innate immune response due to bacterial accumulation in the intestine^148^. Furthermore, it has been shown that the intestinally-produced and secreted innate immunity related protein IRG-7 can lead to the activation of the p38-ATF-7 pathway and is required for the longevity in germlineless nematodes^149^. Apart from long-lived mutants, PMK-1 expression was also observed to decline with normal age, leading to an innate immunosenescence in *C. elegans* that has been proposed to be a driving factor of the aging process^150^. This immunosenescence and the overall involvement of the innate immune system in aging has also been shown in other model organisms^77,151–153^ and might demonstrate an evolutionary conservation. Our work falls in line with these reports and supports an important role of the innate immune response in *C. elegans* aging.

### Neuronal signaling

Our model also shows an enrichment in the neuropeptide signaling pathway. Neuronal communication is important for the maintenance of homeostasis in response to different stressors and a changing environment and has therefore been implicated to have a major role in survival and the aging process^154^. Even in *C. elegans* a cognitive decline can be observed with aging that can be modulated by common longevity pathways like insulin-like signaling^155^. It has also recently been shown that the suppression of excitatory neurotransmitter and neuropeptide signaling is partially required for the longevity of *daf-2* mutants^156^ and similarly a glia-derived neuropeptide signaling pathway that affects the aging rate and healthspan of worms has been described and shows the potential for neuropeptide involvement in the aging process^157^. In line with this, we find *hic-1* and *dop-4* in our predictor gene set. *hic-1* is important for the regulation of acetylcholine neurotransmitter^118^ and might therefore indicate a role of *hic-1* in the locomotion defect that occurs with aging^158^. Besides the role of *dop-4* in the innate immune response^119^, it has also been implicated to slow down habituation^159^. Older worms have been shown to exhibit a greater habituation and a slower recovery from it^160^. The fact that *dop-4* has a negative coefficient in our age prediction suggests that it is less transcribed in older worm populations and thereby making it an interesting target for the cause of increasing habituation with age.

### Human Data

Lastly, we demonstrated that binarized gene expression data also allows building an accurate human age prediction. Currently, the analysis is limited by the data amount and future studies should include more high-quality data from different cohorts with different environments and populations. Optimally, the data would be generated with biopsies from different tissues of living donors without the need of cell culture. Nevertheless, we demonstrated that binarization improves the level of prediction beyond the current standard and that it also allows for a prediction by an elastic net regression, which results in an easy interpretable gene set. Interestingly, we found a significant enrichment in the biological process regulation of cell death, including FOXO1, which indicates that certain age-related pathways, like insulin signaling are indeed relevant for multiple species and evolutionary conserved.

### Conclusions

The binarized expression of our 576 genes is sufficient to predict the biological age of *C. elegans* independent of the underlying genetics or environment with an accuracy near the theoretical limit. Our analysis suggests that the innate immune response, neuronal signaling and single transcription factors are major regulators of the aging process independent of the strain and treatment. Although the temporal rescaling approaches will not be applicable in humans, we have also shown how the binarization approach improves the chronological age prediction of a recent human dataset. Our work establishes that an accurate aging predictor can be built on binarized transcriptomic data that extends beyond the training data, predicts lifespan effects across diverse genetic, environmental or therapeutic interventions, is employable in distinct species and might thus serve as a universally applicable aging clock.

## Supporting information

FileS1_PythonCode

## Acknowledgements

We thank the Regional Computing Center of the University of Cologne (RRZK) for providing computing time on the DFG-funded High Performance Computing (HPC) system CHEOPS as well as support. B.S. acknowledges funding from the Deutsche Forschungsgemeinschaft (SCHU 2494/3-1, SCHU 2494/7-1, SCHU 2494/10-1, SCHU 2494/11-1, CECAD, SFB 829, SFB 670, KFO 286, KFO 329, and GRK2407) and the Deutsche Krebshilfe (70112899).

The authors declare no competing interests.

## Author contributions

D.M. conceived and designed the study and performed all bioinformatics analysis, B.S. coordinated the project and together with D.M. designed the study. All authors wrote the paper.

## Materials and methods

### Data processing

The quality of the data was checked with FastQC^161^ and the data were preprocessed with Fastp^162^ with the following parameters: -g to trim polyG read tails caused by sequencing artifacts, -x to trim polyX, - q 30 for base quality filtering and -e 30 to filter for an average quality score. Paired end samples were processed together. After preprocessing the samples were mapped with STAR-2.7.1a^163^ with the following parameters: --outFilterType BySJout --outFilterMultimapNmax 20 --alignSJoverhangMin 8 -- alignSJDBoverhangMin 1 --outFilterMismatchNmax 999 --outFilterMismatchNoverReadLmax 0.04 -- alignIntronMin 20 --alignIntronMax 1000000 --alignMatesGapMax 1000000 --quantMode GeneCounts.

The genome directories were generated with the ce11 genome, WBcel235.96 without rRNA and the parameter –genomeSAindexNbases 12 for *C. elegans* and the hg38 genome, GRCh38.97 without rRNA and the parameter –genomeSAindexNbases 14 for human data. The parameter –sjdbOverhang was set to the read length of the sample -1.

The counts for unstranded RNA-seq were merged into one .csv file and edgeR^164^ was used to generate Count-Per-Millions (CPM).

### Binarization

To binarize the data first zero CPMs were masked by NaN. For the remaining data the median for each sample was calculated and genes bigger the median were set to 1, while genes smaller or equal to the median were set to 0, finally genes masked by NaN were set to 0 as well.

### Temporal Rescaling

For the temporal rescaling we set the median lifespan of a standard worm to 15.5 days adulthood. We calculated a correction factor for every sample by dividing this standard lifespan by the median lifespan reported by the paper of the corresponding sample. The chronological age of each sample is multiplied with this correction factor to result in the approximated biological age of the sample. The chronological age, correction factor and biological age for every sample can be seen in Table S1.

### 2^nd^ Rescaling Approach

For the 2^nd^ rescaling of the biological age we set the maximum biological age of the worm to 15.5 days. Assuming a normal distribution of biological age around the chronological age of a worm population and further assuming that, on average, worms will die according to their biological age, we can assume that the maximum biological age of a worm is the median lifespan of 15.5 days. Worms living longer than the median lifespan were biologically younger and therefore did not cross the line of 15.5 days. Since the first worms start dying at around 9 days of adulthood, the oldest worms at day 8 should be biologically around 15.5 days old. Therefore, we approximated the standard deviation to be 8/3. Centering a normal distribution at 8 days with a SD of 8/3 will contain 99.73 % of the area under the curve within day 0 to day 16.

Next, we approximated that the biological age distribution is not changing over time and that the SD over 8/3 stays stable. To calculate the median of the data after trimming the data at the maximum age of 15.5 days we first need to calculate how much data is trimmed. We approximate this by utilizing the error function:

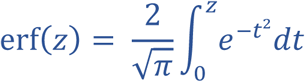

implemented in the Scipy^165^ library.

The approximation of the percentage *p* of data that is remaining on the left side from the maximum lifespan of 15.5 days on the biological age x is:

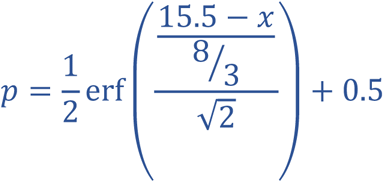

We can divide *p* by 2 to get the approximation of the new median percentage.

To calculate the median in days, we need to revert the calculation. First, we subtract the new number from 0.5 to get the deviation from the original median and use the inverse error function to approximate *s*, the number of standard deviations that the new median is shifted to the left of the old median:

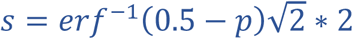

The new median *m*, in other words the new rescaled biological age, can then be calculated by:

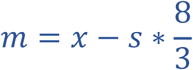

where 8/3 is the standard deviation that we set in the beginning and *x* the biological age, i.e. the original median.

### Model fitting – Parameter search

The age prediction models use an elastic net regression as implemented by Pythons sklearn^166^. The random_state was set to 0, the max_iter to 1000 and positive=False. The best parameter settings for alpha and the L1/L2 ratio were selected using a parameter grid search with a nested cross-validation approach. To avoid overfitting during the training we split the data into multiple partitions. Every sample of the same genetic background, with the same treatment and RNAi interference of the same rounded biological age to days were considered to be one partition. This makes sure, that samples with a similar transcriptome are taken out together during the process. The elastic net regression is trained on the remaining data and the partition that got taken out will be predicted. To get an overview of the accuracy of the model this process is repeated for the partitions in the dataset. In the end every sample will be predicted exactly once, which allows the comparison of the predicted with the true biological age.

A simple cross-validation like this gives an overview of the accuracy of the model, however, to select the best parameter setting a nested cross-validation is required, since otherwise information may leak into the model and introduce another type of overfitting. Therefore, after splitting the data into the test and the train partitions (the outer loop), the latter will be split again into an inner test and train partition (the inner loop). This inner cross-validation will be computed for every parameter-set to compute the average of the absolute error for each parameter setting.

This will be done for every partitioning in the outer loop to select the most stable parameter-set. The parameters selected by this approach for the binarized data are alpha=0.075 and l1_ratio=0.3.

### Model fitting – Optimal Gene Set

To obtain the optimal gene set without overfitting a similar approach was taken. Instead of looping over different parameter settings the cross-validation for the gene set loops over a list of the genes with the highest absolute coefficients. First, for every training partition in the outer loop the full model with alpha=0.075 and l1_ratio=0.3 is computed. This will result in a model, where every gene is annotated with a coefficient. In the binarized model, the sum of the coefficients for all genes that are 1 in the sample equals the predicted age. Therefore, a negative coefficient will result in a younger predicted age, while a positive coefficient will increase the predicted age. Next, we loop over different subsets of the top genes to identify the approximately optimal and smallest gene set for the given partition. For every gene set the inner cross-validation loop is computed and the gene set with the smallest average absolute error is saved. This will be done again for every partition in the outer loop to gain multiple gene sets. Similar to the parameter search the most stable gene set is taken by retaining only those genes that were used by every partition. This stable gene set selected by this approach for the binarized data are the 576 genes described in Table S2.

### Motif Search

The set of genes with a coefficient > 0, respective ≤ 0, were used as input for the findMotifs function of Homer-4.9.1-6^167^ with the parameters -len 8,10 -start 300 -end 100. To make sure that the maximum number of genes got recognized by Homer we first converted the Wormbase IDs to the sequence name with WormBase’s SimpleMine^168^, and added ‘CELE_’ in front of it. These identifiers were then searched in the ‘worm.description’ file of Homer to gain the corresponding RefSeq IDs that are recognized by the program. The p-values were calculated with a hypergeometric test.

### Citation of age predictors from the literature

Because currently no general consensus of quality assessment exists and different measurements are being reported, we state the measurements as reported in the cited paper in the introduction. Some of the most common used assessments are:

1. Mean absolute error (MAE): the mean of the absolute difference of predicted and true age.
2. Root-mean-square-error (RMSE): the square root of the average squared differences. Larger errors have a larger effect on the RMSE than on MAE.
3. Median absolute deviation (MAD): the median absolute difference of predicted and true age.
4. Pearson correlation (r): measurement of how the predicted and true age changes together. Evaluates linear relationships.
5. Spearman correlation (r): similar to Pearson correlation, but evaluates the monotonic relationship. Other than Pearson correlation the variables do not need to change at a linear rate.
6. Coefficient of determination (r^2^): the fraction of the variance that is predictable with the model. Often the r^2^ is the square of the correlation coefficient, however, this is not true in the general case

## Supporting Information Legends

**Figure S1. Alternative models**

Results of the biological age prediction computed by cross-validation. The x-axis shows the rescaled biological age in days starting from adulthood. The y-axis shows the predicted age computed by an elastic net regression on unbinarized CPMs. Every blue dot displays one RNA-seq sample. The regression line is shown in blue and the dotted line shows the perfect linear correlation. The distribution of the data is shown on the side of the plot. MAE= mean absolute error, MAD= median absolute deviation.

**Figure S2. Comparison of the binarized and unbinarized model error**

(A) The absolute error distribution between the predicted and true biological age is plotted for either the unbinarized (red) or binarized (blue) data. The x-axis shows the true biological age in days. The y-axis the absolute error in days. While the unbinarized model strongly increases the absolute prediction error with age, the increase is less pronounced with the binarized model.

(B) The bar plots show the standard deviation of the absolute prediction errors in days. The x-axis shows the true biological age in days. While the binarized model stays relatively stable over age, the unbinarized model increases the variance in the prediction error.

**Figure S3. Explanation of the 2nd rescaling**

(A, C, E) Standard lifespan curves of *C. elegans* with a median lifespan of 15.5 days. The X mark the chronological age for which we show the hypothetical age distributions in (B, D, F) respectively. (B, D, F) show the biological age distribution around the chronological age marked by the X. The biggest portion of the age-synchronized worm population will be as old as the chronological age. However, assuming a normal distribution of the biological age, we can assume that a part of the population is biologically younger, respective older. The green lines indicate the median biological age of the living worm population. The dotted line displays the maximum lifespan.

(A, B) All non-censored worms are still alive in the population, i.e. no worm crossed the maximum lifespan line. The population age median is equal to the peak of the distribution.

(C, D) The first (biologically older) worms died, leading to a truncation of the alive distribution of biological age in the population. This has the consequence that the true median of the alive fraction of the worms will be shifted to the left, away from the peak of the distribution.

(E, F) At the median lifespan, 50 % of the population has died. Assuming a uniform shift of the biological age distribution results in the truncation of the right half of the distribution. The true population median is therefore even further shifted to the left.

**Figure S4. Comparison of the model with random genes and the theoretical limit**

(A) Results of the biological age prediction computed by cross-validation. The x-axis shows the twice rescaled biological age in days starting from adulthood. The y-axis shows the predicted age computed by the elastic net regression after the second rescaling approach. Every blue dot displays one RNA-seq sample. The regression line is shown in blue and the dotted line shows the perfect linear correlation. The distribution of the data is shown on the side of the plot. MAE= mean absolute error, MAD= median absolute deviation.

(B) The y-axis shows the mean absolute error (MAE), respective the median absolute deviation (MAD) of a given prediction in days. The box plots display 1000 random models with 576 genes. The prediction by our final model with a MAE of 0.45 and a MAD of 0.32 is shown as the blue dots and indicated by arrows. The dotted lines show the theoretical limit of prediction given by the limit of accuracy in the chronological age annotation as well as variance in the lifespan data used for rescaling.

**Figure S5. Comparison of our gene set to published gene sets**

Results of the biological age prediction computed by cross-validation based on different gene sets predicted by Tarkhov et al.^97^. The x-axes show the rescaled biological age in days starting from adulthood. The y-axes show the predicted age computed by an elastic net regression on unbinarized (A, B, C) or binarized (D, E, F) gene expression data. Every blue dot displays one RNA-seq sample. The regression lines are shown in blue and the dotted lines show the perfect linear correlation. The distribution of the data is shown on the side of the plot. MAE= mean absolute error, MAD= median absolute deviation.

(A) Prediction based on the unbinarized CPMs of 327 genes generated by a meta-analysis of publicly available microarray data.

(B) Prediction based on the unbinarized CPMs of 902 age-associated genes generated by an RNA-seq experiment.

(C) Prediction based on the unbinarized CPMs of a sparse subset with 71 genes.

(D) Prediction based on the binarized CPMs of 327 genes generated by a meta-analysis of publicly available microarray data.

(E) Prediction based on the binarized CPMs of 902 age-associated genes generated by an RNA-seq experiment.

(F) Prediction based on the binarized CPMs of a sparse subset with 71 genes.

**Figure S6. Chromosome enrichment**

(A) Chromosome distribution of the 286 protein-coding predictor genes with a coefficient <=0 in blue and the number of protein-coding genes that would be expected if the genes were randomly distributed among the chromosomes in red.

(B) Chromosome distribution of the 260 protein-coding predictor genes with a coefficient >0 in blue and the number of protein-coding genes that would be expected if the genes were randomly distributed among the chromosomes in red.

(C) Differences of the observed to the expected numbers in percent for the protein-coding genes with a coefficient >0 in blue and with a coefficient <=0 in red.

*p<0.05, **p<=0.01, ***p<=0.001, Hypergeometric tests were performed and the resulting p-values were corrected with the Benjamini-Hochberg procedure. Table S4 contains more detailed statistics.

**Figure S7. Motif enrichment nearby the TSS**

Results of a motif enrichment analysis for the region −300 bp to +100 bp from the transcription start site of the genes with a coefficient <=0 (A) and genes with a coefficient >0 (B). The columns show the name of the transcription factor in the first column with the known motif in the second column. Column 3 and 4 show the percentage of target genes, respective background genes, containing the motif in the described region. Column 5 shows the fold change enrichment, column 6 the corresponding Hypergeometric p-value and the last column the Benjamini-Hochberg adjusted q-value.

**Figure S8. Unbinarized human data**

(A) Results of the age prediction computed by cross-validation on human fibroblast gene expression data. The x-axis shows the chronological age in years. The y-axis shows the predicted age computed by an elastic net regression on unbinarized gene expression data. Every blue dot displays one RNA-seq sample. The regression line is shown in blue and the dotted line shows the perfect linear correlation. The distribution of the data is shown on the side of the plot. MAE= mean absolute error, MAD= median absolute deviation.

(B) Box plots of age predictions of samples from Hutchinson–Gilford progeria syndrome patients (red) and predictions of age-matched healthy controls (blue) by the elastic net regression of unbinarized gene expression data. Progeria samples show no significant increase in the predicted age compared to age-matched healthy controls.

The p-value was calculated by an independent t-test. Table S4 contains more detailed statistics.

**Table S1. Data overview**

**Table S2. *C. elegans* age prediction gene set Table S3. Lifespan variation**

**Table S4. Statistics**

**Table S5. Human age prediction gene set**

**File S1. Python code**

## Notes

### Competing Interest Statement

The authors have declared no competing interest.

## References

1. López-Otín, C., Blasco, M. A., Partridge, L., Serrano, M. & Kroemer, G. The Hallmarks of Aging. Cell 153, 1194–1217 (2013).

2. Belsky, D. W. et al. Change in the Rate of Biological Aging in Response to Caloric Restriction: CALERIE Biobank Analysis. Journals Gerontol. - Ser. A Biol. Sci. Med. Sci. 73, 4–10 (2018).

3. Stubbs, T. M. et al. Multi-tissue DNA methylation age predictor in mouse. 1–14 (2017). doi: 10.1186/s13059-017-1203-5

4. Horvath, S. et al. Reversing age: dual species measurement of epigenetic age with a single clock. bioRxiv (2020).

5. Lujan, C. et al. A CellAgeClock for expedited discovery of anti-ageing compounds. bioRxiv (2020).

6. Chen, B. H. et al. DNA methylation - based measures of biological age: meta - analysis predicting time to death. Aging (Albany. NY). 8, 1844–1859 (2016).

7. Jylhävä, J., Pedersen, N. L. & Hägg, S. Biological Age Predictors. EBioMedicine 21, 29–36 (2017).

8. Galkin, F. et al. Biohorology and biomarkers of aging: current state-of-the-art, challenges and opportunities. Ageing Res. Rev. 101050 (2020). doi: 10.1016/j.arr.2020.101050

9. Xia, X., Chen, W., Mcdermott, J. & Han, J. J. Molecular and phenotypic biomarkers of aging. F1000Research 6, (2017).

10. Debès, C., Leote, A. C. & Beyer, A. Computational approaches for the systematic analysis of aging-associated molecular alterations. Drug Discovery Today: Disease Models 27, 51–59 (2018).

11. Zhavoronkov, A. & Mamoshina, P. Deep Aging Clocks: The Emergence of AI-Based Biomarkers of Aging and Longevity. Trends Pharmacol. Sci. 40, 546–549 (2019).

12. Kane, A. et al. MACHINE LEARNING ANALYSIS OF MOUSE FRAILTY FOR PREDICTION OF BIOLOGICAL AGE AND LIFE EXPECTANCY. Innov. Aging 3, S903–S903 (2019).

13. Zhavoronkov, A. et al. Artificial intelligence for aging and longevity research: Recent advances and perspectives. Ageing Res. Rev. 49, 49–66 (2019).

14. Kudryashova, K. S., Burka, K., Kulaga, A. Y., Vorobyeva, N. S. & Kennedy, B. K. Aging Biomarkers: From Functional Tests to Multi-Omics Approaches. Proteomics 1900408, 1–15 (2020).

15. Ferrucci, L. et al. Measuring biological aging in humans: A quest. Aging Cell 19, 1–21 (2020).

16. Solovev, I. Multi-omics approaches to human biological age estimation. Mech. Ageing Dev. 185, 111192 (2020).

17. Putin, E. et al. Deep biomarkers of human aging: Application of deep neural networks to biomarker development. Aging (Albany. NY). 8, 1021–1030 (2016).

18. Wang, Z. et al. Predicting age by mining electronic medical records with deep learning characterizes di ff erences between chronological and physiological age. 76, 59–68 (2017).

19. Mamoshina, P. et al. Population specific biomarkers of human aging: A big data study using South Korean, Canadian, and Eastern European patient populations. Journals Gerontol. - Ser. A Biol. Sci. Med. Sci. 73, 1482–1490 (2018).

20. Pyrkov, T. V. et al. Extracting biological age from biomedical data via deep learning: Too much of a good thing? Sci. Rep. 8, 1–11 (2018).

21. Zhong, X. et al. Estimating Biological Age in the Singapore Longitudinal Aging Study. Journals Gerontol. Ser. A XX, 1–8 (2019).

22. Hannum, G. et al. Genome-wide Methylation Profiles Reveal Quantitative Views of Human Aging Rates. Mol. Cell 49, 359–367 (2013).

23. Horvath, S. DNA methylation age of human tissues and cell types. Genome Biol. 16, 96 (2015).

24. Horvath, S. & Raj, K. DNA methylation-based biomarkers and the epigenetic clock theory of ageing. Nat. Rev. Genet. 19, 371–384 (2018).

25. Field, A. E. et al. DNA Methylation Clocks in Aging: Categories, Causes, and Consequences. Mol. Cell 71, 882–895 (2018).

26. Vidal-bralo, L., Lopez-golan, Y. & Gonzalez, A. Simplified Assay for Epigenetic Age Estimation in Whole Blood of Adults. 7, 1–7 (2016).

27. Thompson, M. J. et al. A multi-tissue full lifespan epigenetic clock for mice. Aging (Albany. NY). 10, 2832–2854 (2018).

28. Bell, C. G. et al. DNA methylation aging clocks: challenges and recommendations. Genome Biol. 20, 249 (2019).

29. Zhang, Q. et al. Improved prediction of chronological age from DNA methylation limits it as a biomarker of ageing. bioRxiv 327890 (2018). doi: 10.1101/327890

30. Hernando-Herraez, I. et al. Ageing affects DNA methylation drift and transcriptional cell-to-cell variability in mouse muscle stem cells. Nat. Commun. 10, 1–11 (2019).

31. Michalak, E. M., Burr, M. L., Bannister, A. J. & Dawson, M. A. The roles of DNA, RNA and histone methylation in ageing and cancer. Nat. Rev. Mol. Cell Biol. 20, 573–589 (2019).

32. Feser, J. et al. Elevated Histone Expression Promotes Life Span Extension. Mol. Cell 39, 724–735 (2010).

33. Greer, E. L. et al. Transgenerational epigenetic inheritance of longevity in Caenorhabditis elegans. Nature 479, 365–371 (2011).

34. Maures, T. J., Greer, E. L., Hauswirth, A. G. & Brunet, A. The H3K27 demethylase UTX-1 regulates C. elegans lifespan in a germline-independent, insulin-dependent manner. Aging Cell 10, 980–990 (2011).

35. Cao, X. & Dang, W. Histone Modification Changes During Aging. in Epigenetics of Aging and Longevity 309–328 (Elsevier, 2018). doi: 10.1016/B978-0-12-811060-7.00015-2

36. Wang, Y., Yuan, Q. & Xie, L. Histone Modifications in Aging: The Underlying Mechanisms and Implications. Curr. Stem Cell Res. Ther. 13, 125–135 (2018).

37. Feser, J. & Tyler, J. Chromatin structure as a mediator of aging. FEBS Lett. 585, 2041–2048 (2011).

38. Criscione, S. W., Teo, Y. V. & Neretti, N. The Chromatin Landscape of Cellular Senescence. Trends Genet. 32, 751–761 (2016).

39. Sun, L., Yu, R. & Dang, W. Chromatin architectural changes during cellular senescence and aging. Genes (Basel). 9, (2018).

40. Bahar, R. et al. Increased cell-to-cell variation in gene expression in ageing mouse heart. Nature 441, 1011–1014 (2006).

41. Southworth, L. K., Owen, A. B. & Kim, S. K. Aging mice show a decreasing correlation of gene expression within genetic modules. PLoS Genet. 5, (2009).

42. Viñuela, A., Snoek, L. B., Riksen, J. A. G. & Kammenga, J. E. Genome-wide gene expression regulation as a function of genotype and age in C. elegans. Genome Res. 20, 929–937 (2010).

43. Kogan, V., Molodtsov, I., Menshikov, L. I., Reis, R. J. S. & Fedichev, P. Stability analysis of a model gene network links aging, stress resistance, and negligible senescence. Sci. Rep. 5, 1–12 (2015).

44. Stegeman, R. & Weake, V. M. Transcriptional Signatures of Aging. J. Mol. Biol. 429, 2427–2437 (2017).

45. Bryois, J. et al. Time-dependent genetic effects on gene expression implicate aging processes. Genome Res. 27, 545–552 (2017).

46. Lai, R. W. et al. Multi-level remodeling of transcriptional landscapes in aging and longevity. BMB Rep. 52, 86–108 (2019).

47. Debès, C. et al. Aging-associated changes in transcriptional elongation influence metazoan longevity. bioRxiv 719864 (2019). doi: 10.1101/719864

48. Stoeger, T. et al. Aging is associated with a systemic length-driven transcriptome imbalance. bioRxiv 2, 691154 (2019).

49. Frenk, S. & Houseley, J. Gene expression hallmarks of cellular ageing. Biogerontology 19, 547–566 (2018).

50. Rangaraju, S. et al. Suppression of transcriptional drift extends C. elegans lifespan by postponing the onset of mortality. Elife 4, e08833 (2015).

51. Sood, S. et al. A novel multi-tissue RNA diagnostic of healthy ageing relates to cognitive health status. Genome Biol. 1–17 (2015). doi: 10.1186/s13059-015-0750-x

52. Jacob, L. & Speed, T. P. The healthy ageing gene expression signature for Alzheimer’s disease diagnosis: a random sampling perspective. Genome Biol. 19, 97 (2018).

53. Timmons, J. A. et al. A statistical and biological response to an informatics appraisal of healthy aging gene signatures. Genome Biol. 20, 4–7 (2019).

54. Pilling, L. C. et al. The reported healthy ageing gene expression score: lack of association in two cohorts. bioRxiv 44, 2–7 (2015).

55. Janssens, G. E. et al. Transcriptomics-Based Screening Identifies Pharmacological Inhibition of Hsp90 as a Means to Defer Aging. Cell Rep. 27, 467-480.e6 (2019).

56. Peters, M. J. et al. The transcriptional landscape of age in human peripheral blood. Nat. Commun. 6, 8570 (2015).

57. Mamoshina, P. et al. Machine learning on human muscle transcriptomic data for biomarker discovery and tissue-specific drug target identification. Front. Genet. 9, 1–10 (2018).

58. González-Velasco, O., Papy-García, D., Le Douaron, G., Sánchez-Santos, J. M. & De Las Rivas, J. Transcriptomic landscape, gene signatures and regulatory profile of aging in the human brain. Biochim. Biophys. Acta - Gene Regul. Mech. 194491 (2020). doi: 10.1016/j.bbagrm.2020.194491

59. Carithers, L. J. et al. A Novel Approach to High-Quality Postmortem Tissue Procurement: The GTEx Project. Biopreserv. Biobank. 13, 311–317 (2015).

60. Yang, J., Huang, T., Petralia, F., Long, Q. & Zhang, B. Synchronized age-related gene expression changes across multiple tissues in human and the link to complex diseases. Sci. Rep. 1–16 (2015). doi: 10.1038/srep15145

61. Fleischer, J. G. et al. Predicting age from the transcriptome of human dermal fibroblasts. Genome Biol. 19, 1–8 (2018).

62. Wang, K. et al. Comprehensive map of age-associated splicing changes across human tissues and their contributions to age-associated diseases. Sci. Rep. 8, 1–12 (2018).

63. Ren, X. & Kuan, P. F. RNAAgeCalc: A multi-tissue transcriptional age calculator. bioRxiv 2020.02.14.950188 (2020). doi: 10.1101/2020.02.14.950188

64. Huan, T. et al. Age-associated microRNA expression in human peripheral blood is associated with all-cause mortality and age-related traits. Aging Cell 1–10 (2018). doi: 10.1111/acel.12687

65. Choukrallah, M., Hoeng, J., Peitsch, M. C. & Martin, F. Lung transcriptomic clock predicts premature aging in cigarette smoke-exposed mice. BMC Genomics 1–9 (2020).

66. Tanaka, T. et al. Plasma proteomic signature of age in healthy humans. Aging Cell 17, 1–13 (2018).

67. Lehallier, B. et al. Undulating changes in human plasma proteome profiles across the lifespan. Nat. Med. 25, 1843–1850 (2019).

68. Earls, J. C. et al. Multi-Omic Biological Age Estimation and Its Correlation With Wellness and Disease Phenotypes: A Longitudinal Study of 3, 558 Individuals. Journals Gerontol. - Ser. A Biol. Sci. Med. Sci. 74, 52–60 (2019).

69. Johnson, A. A., Shokhirev, M. N., Wyss-Coray, T. & Lehallier, B. Systematic review and analysis of human proteomics aging studies unveils a novel proteomic aging clock and identifies key processes that change with age. Agein Res. Rev. 116544 (2020). doi: 10.1016/j.jns.2019.116544

70. Liu, Z. et al. A new aging measure captures morbidity and mortality risk across diverse subpopulations from NHANES IV: A cohort study. PLoS Med. 16, e1002760 (2019).

71. Zhou, W. et al. Longitudinal multi-omics of host–microbe dynamics in prediabetes. Nature 569, 663–671 (2019).

72. Schüssler-Fiorenza Rose, S. M. et al. A longitudinal big data approach for precision health. Nat. Med. 25, 792–804 (2019).

73. Ahadi, S. et al. Personal aging markers and ageotypes revealed by deep longitudinal profiling. Nat. Med. 26, 83–90 (2020).

74. Balliu, B. et al. Genetic regulation of gene expression and splicing during a 10-year period of human aging. Genome Biol. 20, 1–16 (2019).

75. Brown, A. A. et al. Age-dependent changes in mean and variance of gene expression across tissues in a twin cohort. Hum. Mol. Genet. 27, 732–741 (2018).

76. Horvath, S. et al. An epigenetic clock analysis of race/ethnicity, sex, and coronary heart disease. Genome Biol. 17, 0–22 (2016).

77. Benayoun, B. A. et al. Remodeling of epigenome and transcriptome landscapes with aging in mice reveals widespread induction of inflammatory responses. Genome Res. 29, 697–709 (2019).

78. Yu, Y. et al. A rat RNA-Seq transcriptomic BodyMap across 11 organs and 4 developmental stages. Nat. Commun. 5, 3230 (2014).

79. Shavlakadze, T. et al. Age-Related Gene Expression Signature in Rats Demonstrate Early, Late, and Linear Transcriptional Changes from Multiple Tissues Resource Age-Related Gene Expression Signature in Rats Demonstrate Early, Late, and Linear Transcriptional Changes from Multipl. Cell Rep. 28, 3263–3273 (2019).

80. Kim, S. K. Common aging pathways in worms, flies, mice and humans. J. Exp. Biol. 210, 1607–1612 (2007).

81. de Magalhães, J. P., Curado, J. & Church, G. M. Meta-analysis of age-related gene expression profiles identifies common signatures of aging. Bioinformatics 25, 875–881 (2009).

82. Palmer, D., Fabris, F., Doherty, A., Freitas, A. A. & Magalhães, J. P. de. Ageing Transcriptome Meta-Analysis Reveals Similarities Between Key Mammalian Tissues. bioRxiv 815381 (2019). doi: 10.1101/815381

83. Johnson, T. E. Aging can be genetically dissected into component processes using long-lived lines of Caenorhabditis elegans. Proc. Natl. Acad. Sci. U. S. A. 84, 3777–3781 (1987).

84. Johnson, T. E. & Lithgow, G. J. The Search for the Genetic Basis of Aging: The Identification of Gerontogenes in the Nematode Caenorhabditis elegans. J. Am. Geriatr. Soc. 40, 936–945 (1992).

85. Gershon, H. & Gershon, D. Caenorhabditis elegans - A paradigm for aging research: Advantages and limitations. Mech. Ageing Dev. 123, 261–274 (2002).

86. Johnson, T. E. Advantages and disadvantages of Caenorhabditis elegans for aging research. Exp. Gerontol. 38, 1329–1332 (2003).

87. Henderson, S. T., Rea, S. L. & Johnson, T. E. Dissecting the process of aging using the nematode Caenorhabditis elegans. Handb. Biol. aging 360–399 (2005).

88. Pincus, Z. & Slack, F. J. Developmental biomarkers of aging in Caenorhabditis elegans. Dev. Dyn. 239, 1306–1314 (2010).

89. Rea, S. L., Wu, D., Cypser, J. R., Vaupel, J. W. & Johnson, T. E. A stress-sensitive reporter predicts longevity in isogenic populations of Caenorhabditis elegans. Nat. Genet. 37, 894–898 (2005).

90. Cypser, J. R. et al. Predicting longevity in C. elegans: Fertility, mobility and gene expression. Mech. Ageing Dev. 134, 291–297 (2013).

91. Sánchez-Blanco, A. & Kim, S. K. Variable pathogenicity determines individual lifespan in caenorhabditis elegans. PLoS Genet. (2011). doi: 10.1371/journal.pgen.1002047

92. Pincus, Z., Smith-Vikos, T. & Slack, F. J. MicroRNA predictors of longevity in caenorhabditis elegans. PLoS Genet. 7, (2011).

93. Huang, C., Xiong, C. & Kornfeld, K. Measurements of age-related changes of physiological processes that predict lifespan of Caenorhabditis elegans. Proc. Natl. Acad. Sci. U. S. A. 101, 8084–8089 (2004).

94. Martineau, C. N., Brown, A. E. X. & Laurent, P. Multidimensional phenotyping predicts lifespan and quantifies health in C. elegans. bioRxiv 681197 (2019). doi: 10.1101/681197

95. Golden, T. R., Hubbard, A., Dando, C., Herren, M. A. & Melov, S. Age-related behaviors have distinct transcriptional profiles in Caenorhabditis.elegans. Aging Cell 7, 850–865 (2008).

96. Fortney, K., Kotlyar, M. & Jurisica, I. Inferring the functions of longevity genes with modular subnetwork biomarkers of Caenorhabditis elegans aging. Genome Biol. 11, (2010).

97. Tarkhov, A. E. et al. A universal transcriptomic signature of age reveals the temporal scaling of Caenorhabditis elegans aging trajectories. Sci. Rep. 9, 1–18 (2019).

98. Stroustrup, N. et al. The temporal scaling of Caenorhabditis elegans ageing. Nature 530, 103–107 (2016).

99. Hastings, J. et al. Multi-omics and genome-scale modeling reveal a metabolic shift during C. elegans aging. Front. Mol. Biosci. 6, 1–18 (2019).

100. Mann, F. G., Van Nostrand, E. L., Friedland, A. E., Liu, X. & Kim, S. K. Deactivation of the GATA Transcription Factor ELT-2 Is a Major Driver of Normal Aging in C. elegans. PLoS Genet. 12, e1005956.-26 (2016).

101. Petrascheck, M. & Miller, D. L. Computational analysis of lifespan experiment reproducibility. Front. Genet. 8, 1–11 (2017).

102. Zarse, K. et al. Impaired insulin/IGF1 signaling extends life span by promoting mitochondrial L-proline catabolism to induce a transient ROS signal. Cell Metab. 15, 451–465 (2012).

103. Rollins, J. A., Shaffer, D., Snow, S. S., Kapahi, P. & Rogers, A. N. Dietary restriction induces posttranscriptional regulation of longevity genes. Life Sci. Alliance 2, 1–22 (2019).

104. Inukai, S., Pincus, Z., De Lencastre, A. & Slack, F. J. A microRNA feedback loop regulates global microRNA abundance during aging. Rna 24, 159–172 (2018).

105. Nhan, J. D. et al. Redirection of SKN-1 abates the negative metabolic outcomes of a perceived pathogen infection. Proc. Natl. Acad. Sci. 116, 22322–22330 (2019).

106. Nedialkova, D. D. & Leidel, S. A. Optimization of Codon Translation Rates via tRNA Modifications Maintains Proteome Integrity. Cell 161, 1606–1618 (2015).

107. Wu, Z. et al. Dietary Restriction Extends Lifespan through Metabolic Regulation of Innate Immunity. Cell Metab. 29, 1–23 (2019).

108. Sonowal, R. et al. Indoles from commensal bacteria extend healthspan. Proc. Natl. Acad. Sci. U. S. A. 114, E7506–E7515 (2017).

109. Pryor, R. et al. Host-Microbe-Drug-Nutrient Screen Identifies Bacterial Effectors of Metformin Therapy. Cell 178, 1299-1312.e29 (2019).

110. Admasu, T. D. et al. Drug Synergy Slows Aging and Improves Healthspan through IGF and SREBP Lipid Signaling. Dev. Cell 47, 67-79.e5 (2018).

111. Finger, F. et al. Olfaction regulates organismal proteostasis and longevity via microRNA-dependent signalling. Nat. Metab. 1, 350–359 (2019).

112. Vilchez, D. et al. RPN-6 determines C. elegans longevity under proteotoxic stress conditions. Nature 489, 263–268 (2012).

113. Webster, C. M. et al. Genome-wide RNAi Screen for Fat Regulatory Genes in C. elegans Identifies a Proteostasis-AMPK Axis Critical for Starvation Survival. Cell Rep. 20, 627–640 (2017).

114. Yang, W., Dierking, K. & Schulenburg, H. WormExp: A web-based application for a Caenorhabditis elegans-specific gene expression enrichment analysis. Bioinformatics 32, 943–945 (2016).

115. Budovskaya, Y. V. et al. An elt-3/elt-5/elt-6 GATA Transcription Circuit Guides Aging in C. elegans. Cell 134, 291–303 (2008).

116. Szklarczyk, D. et al. STRING v11: Protein-protein association networks with increased coverage, supporting functional discovery in genome-wide experimental datasets. Nucleic Acids Res. 47, D607–D613 (2019).

117. Subhash, S. & Kanduri, C. GeneSCF: A real-time based functional enrichment tool with support for multiple organisms. BMC Bioinformatics 17, 1–10 (2016).

118. Tikiyani, V. et al. Wnt Secretion Is Regulated by the Tetraspan Protein HIC-1 through Its Interaction with Neurabin/NAB-1. Cell Rep. 25, 1856-1871.e6 (2018).

119. Joshi, K. K., Matlack, T. L. & Rongo, C. Dopamine signaling promotes the xenobiotic stress response and protein homeostasis. EMBO J. 35, e201592524.-17 (2016).

120. Martins, R., Lithgow, G. J. & Link, W. Long live FOXO: Unraveling the role of FOXO proteins in aging and longevity. Aging Cell 15, 196–207 (2016).

121. Melzer, D., Pilling, L. C. & Ferrucci, L. The genetics of human ageing. Nat. Rev. Genet. 21, 88–101 (2020).

122. Sen, P., Shah, P. P., Nativio, R. & Berger, S. L. Epigenetic Mechanisms of Longevity and Aging. Cell 166, 822–839 (2016).

123. Verjan, J. C. G., Martinez, E. R. V., Segura, N. A. R. & Campos, R. H. M. The RNA world of human ageing. Hum. Genet. 137, 865–879 (2018).

124. Srivastava, S. Emerging insights into the metabolic alterations in aging using metabolomics. Metabolites 9, 1–16 (2019).

125. Denzel, M. S., Lapierre, L. R. & Mack, H. I. D. Emerging topics in C. elegans aging research: Transcriptional regulation, stress response and epigenetics. Mech. Ageing Dev. 177, 4–21 (2019).

126. Son, H. G., Altintas, O., Kim, E. J. E., Kwon, S. & Lee, S. J. V. Age-dependent changes and biomarkers of aging in Caenorhabditis elegans. Aging Cell 18, 1–11 (2019).

127. Zhao, Y. et al. Two forms of death in ageing Caenorhabditis elegans. Nat. Commun. 8, 1–8 (2017).

128. Nikopoulou, C., Parekh, S. & Tessarz, P. Ageing and sources of transcriptional heterogeneity. Biol. Chem. 400, 867–878 (2019).

129. Pincus, Z. & Slack, F. J. Transcriptional (dys)regulation and aging in Caenorhabditis elegans. Genome Biol. 9, 233 (2008).

130. Enge, M. et al. Single-Cell Analysis of Human Pancreas Reveals Transcriptional Signatures of Aging and Somatic Mutation Patterns. Cell 171, 321-330.e14 (2017).

131. Swisa, A., Kaestner, K. H. & Dor, Y. Transcriptional Noise and Somatic Mutations in the Aging Pancreas. Cell Metab. 26, 809–811 (2017).

132. Liu, P., Song, R., Elison, G. L., Peng, W. & Acar, M. Noise reduction as an emergent property of single-cell aging. Nat. Commun. 8, 1–13 (2017).

133. Lin, X.-X. et al. DAF-16/FOXO and HLH-30/TFEB function as combinatorial transcription factors to promote stress resistance and longevity. Nat. Commun. 1–15 (2018).

134. Senchuk, M. M. et al. Activation of DAF-16/FOXO by reactive oxygen species contributes to longevity in long-lived mitochondrial mutants in Caenorhabditis elegans. PLoS Genet. 14, 1–27 (2018).

135. Sun, X., Chen, W. D. & Wang, Y. D. DAF-16/FOXO transcription factor in aging and longevity. Front. Pharmacol. 8, 1–8 (2017).

136. Bianco, J. N. & Schumacher, B. MPK-1/ERK pathway regulates DNA damage response during development through DAF-16/FOXO. Nucleic Acids Res. 46, 6129–6139 (2018).

137. Li, S. T. et al. DAF-16 stabilizes the aging transcriptome and is activated in mid-aged Caenorhabditis elegans to cope with internal stress. Aging Cell 18, (2019).

138. Block, D. H. & Shapira, M. GATA transcription factors as tissue-specific master regulators for induced responses. Worm 4, e1118607 (2015).

139. Mueller, M. M. et al. DAF-16/FOXO and EGL-27/GATA promote developmental growth in response to persistent somatic DNA damage. Nat. Cell Biol. 16, 1168–1179 (2014).

140. Tepper, R. G. et al. PQM-1 Complements DAF-16 as a Key Transcriptional Regulator of DAF-2-Mediated Development and Longevity. Cell 154, 676–690 (2013).

141. Templeman, N. M. et al. Insulin Signaling Regulates Oocyte Quality Maintenance with Age via Cathepsin B Activity. Curr. Biol. 28, 753-760.e4 (2018).

142. O\textquoterightBrien, D. et al. A PQM-1-Mediated Response Triggers Transcellular Chaperone Signaling and Regulates Organismal Proteostasis. Cell Rep. 23, 3905–3919 (2018).

143. Kurz, C. L. & Tan, M. W. Regulation of aging and innate immunity in C. elegans. Aging Cell 3, 185–193 (2004).

144. Ermolaeva, M. A. & Schumacher, B. Insights from the worm: The C. elegans model for innate immunity. Seminars in Immunology 26, 303–309 (2014).

145. Ermolaeva, M. A. et al. DNA damage in germ cells induces an innate immune response that triggers systemic stress resistance. Nature 501, 416–420 (2013).

146. Schmeisser, S. et al. Neuronal ROS signaling rather than AMPK/sirtuin-mediated energy sensing links dietary restriction to lifespan extension. Mol. Metab. 2, 92–102 (2013).

147. Troemel, E. R. et al. p38 MAPK regulates expression of immune response genes and contributes to longevity in C. elegans. PLoS Genet. 2, e183 (2006).

148. Kumar, S. et al. Lifespan Extension in C. elegans Caused by Bacterial Colonization of the Intestine and Subsequent Activation of an Innate Immune Response. Dev. Cell 49, 100-117.e6 (2019).

149. Yunger, E., Safra, M., Levi-Ferber, M., Haviv-Chesner, A. & Henis-Korenblit, S. Innate immunity mediated longevity and longevity induced by germ cell removal converge on the C-type lectin domain protein IRG-7. PLoS Genet. 13, e1006577 (2017).

150. Youngman, M. J., Rogers, Z. N. & Kim, D. H. A decline in p38 MAPK signaling underlies immunosenescence in Caenorhabditis elegans. PLoS Genet. 7, e1002082 (2011).

151. Müller, L., Fülöp, T. & Pawelec, G. Immunosenescence in vertebrates and invertebrates. Immun. Ageing 10, (2013).

152. Xia, J., Gravato-Nobre, M. & Ligoxygakis, P. Convergence of longevity and immunity: lessons from animal models. Biogerontology 20, 271–278 (2019).

153. Du, L. et al. Transcriptome analysis reveals immune - related gene expression changes with age in giant panda (Ailuropoda melanoleuca) blood. Aging (Albany. NY). 11, (2019).

154. Alcedo, J., Flatt, T. & Pasyukova, E. G. Neuronal inputs and outputs of aging and longevity. Front. Genet. 4, 1–14 (2013).

155. Stein, G. M. & Murphy, C. T. The intersection of aging, longevity pathways, and learning and memory in C. elegans. Front. Genet. 3, 1–13 (2012).

156. Zullo, J. M. et al. Regulation of lifespan by neural excitation and REST. Nature 574, 359–364 (2019).

157. Yin, J. A. et al. Genetic variation in glia-neuron signalling modulates ageing rate. Nature 551, 198–203 (2017).

158. Glenn, C. F. et al. Behavioral deficits during early stages of aging in Caenorhabditis elegans result from locomotory deficits possibly linked to muscle frailty. Journals Gerontol. - Ser. A Biol. Sci. Med. Sci. 59, 1251–1260 (2004).

159. Ardiel, E. L. et al. Dopamine receptor DOP-4 modulates habituation to repetitive photoactivation of a C. elegans polymodal nociceptor. Learn. Mem. 23, 495–503 (2016).

160. Beck, C. D. O. & Rankin, C. H. Effects of aging on habituation in the nematode Caenorhabditis elegans. Behav. Processes 28, 145–163 (1993).

161. Andrews, S., Krueger, F., Segonds-Pichon, Anne Biggins, L., Krueger, C. & Wingett, S. FastQC. (2010).

162. Chen, S., Zhou, Y., Chen, Y. & Gu, J. Fastp: An ultra-fast all-in-one FASTQ preprocessor. Bioinformatics 34, i884–i890 (2018).

163. Dobin, A. et al. STAR: Ultrafast universal RNA-seq aligner. Bioinformatics 29, 15–21 (2013).

164. Robinson, M. D., McCarthy, D. J. & Smyth, G. K. edgeR: A Bioconductor package for differential expression analysis of digital gene expression data. Bioinformatics 26, 139–140 (2009).

165. Virtanen, P. et al. SciPy 1.0: fundamental algorithms for scientific computing in Python. Nat. Methods 17, 261–272 (2020).

166. Varoquaux, G. et al. Scikit-learn: Machine Learning in Python Fabian. J. Mach. Learn. Res. (2011). doi: 10.1145/2786984.2786995

167. Heinz, S. et al. Simple Combinations of Lineage-Determining Transcription Factors Prime cis-Regulatory Elements Required for Macrophage and B Cell Identities. Mol. Cell 38, 576–589 (2010).

168. Harris, T. W. et al. WormBase: a modern Model Organism Information Resource. Nucleic Acids Res. 48, D762–D767 (2020).

169. Sen, P. et al. H3K36 methylation promotes longevity by enhancing transcriptional fidelity. Genes Dev. 29, 1362–1376 (2015).

170. Lan, J. et al. Translational Regulation of Non-autonomous Mitochondrial Stress Response Promotes Longevity. Cell Rep. 1050–1062 (2019). doi: 10.1016/j.celrep.2019.06.078

171. Sellegounder, D. et al. Neuronal GPCR NPR-8 regulates C. elegans defense against pathogen infection. Sci. Adv. 5, 1–16 (2019).

172. Visvikis, O. et al. Innate host defense requires TFEB-mediated transcription of cytoprotective and antimicrobial genes. Immunity 40, 896–909 (2014).

