## Supplementary Figure 1-8 for "A transcriptome based aging clock near the theoretical limit of accuracy"

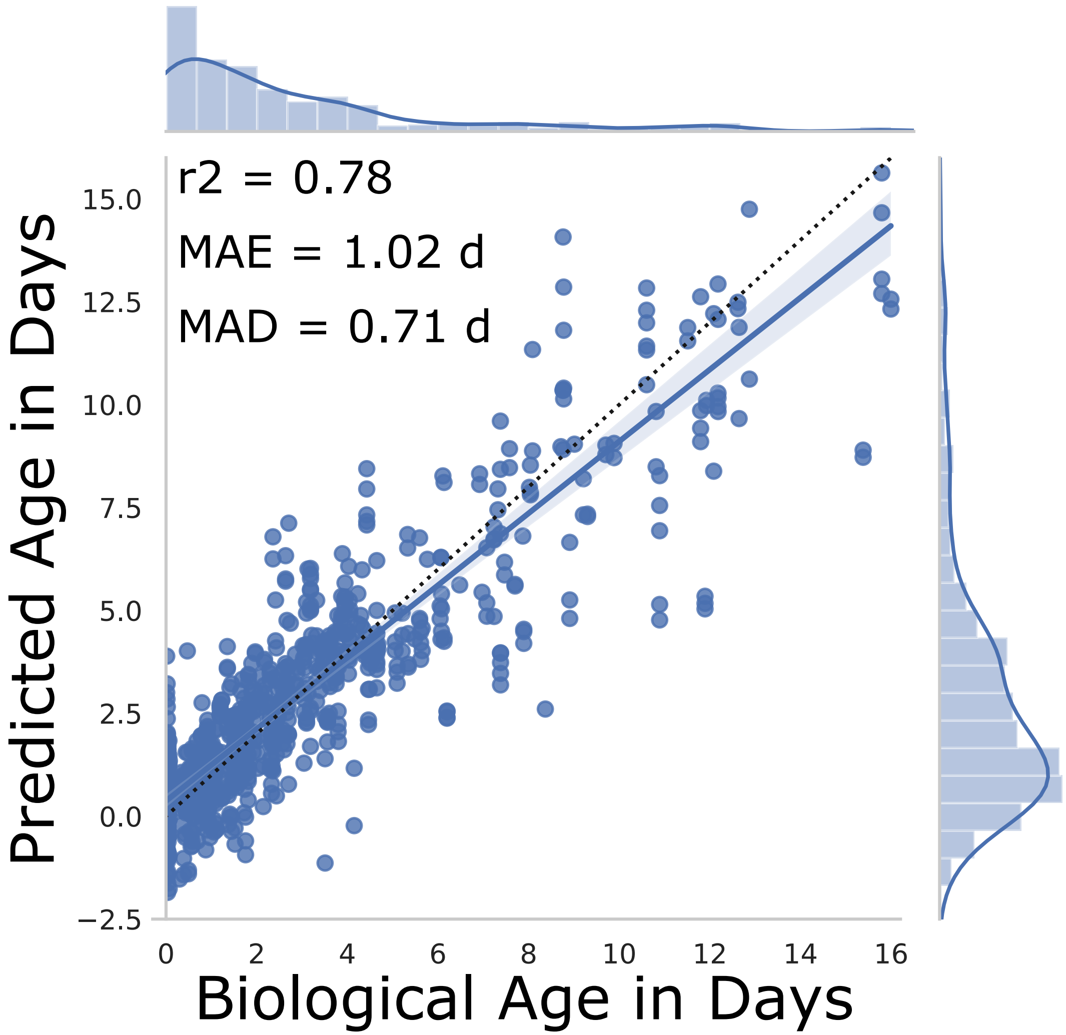

**Fig. S1** **Meyer and Schumacher**

A

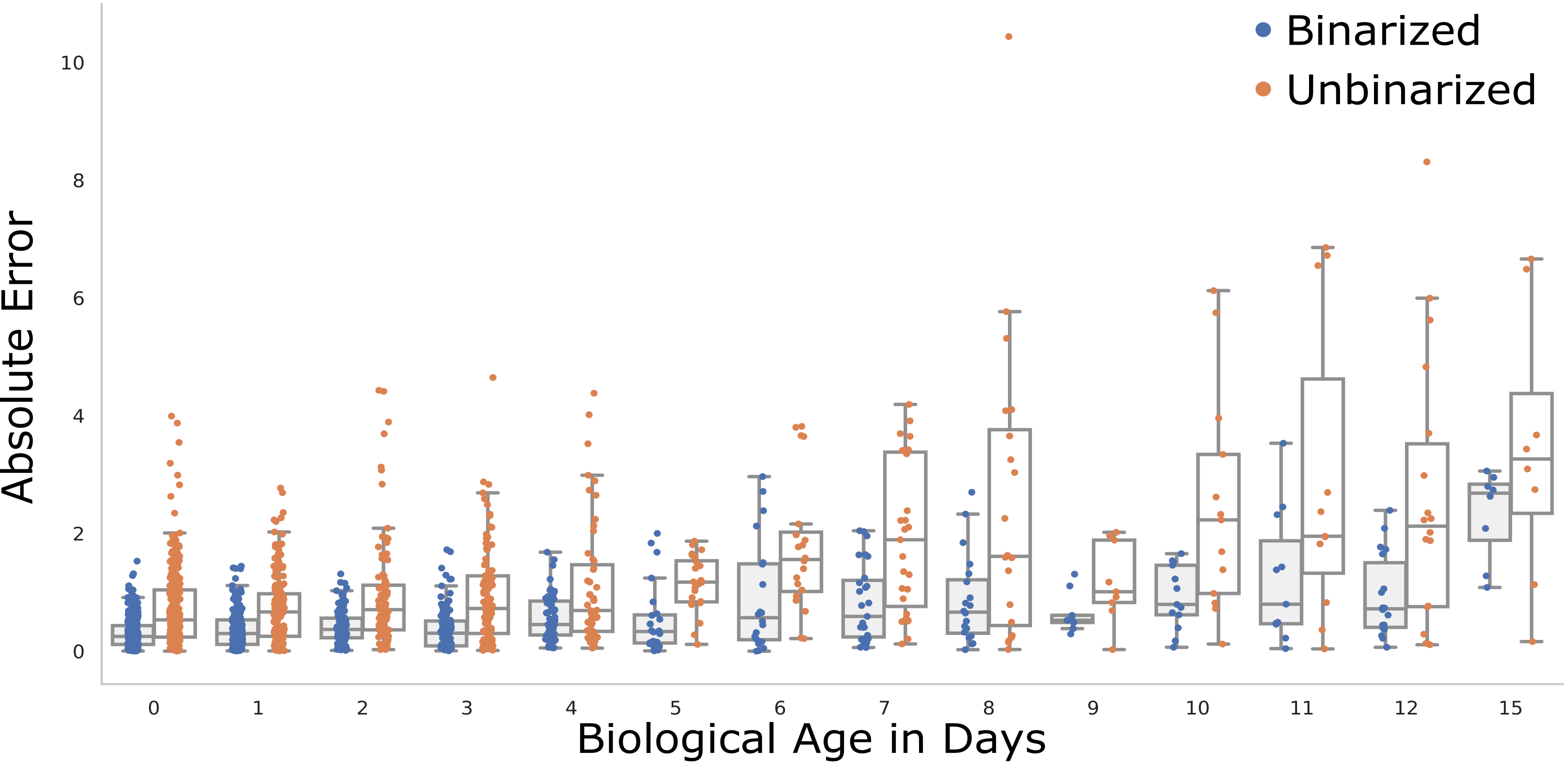

B

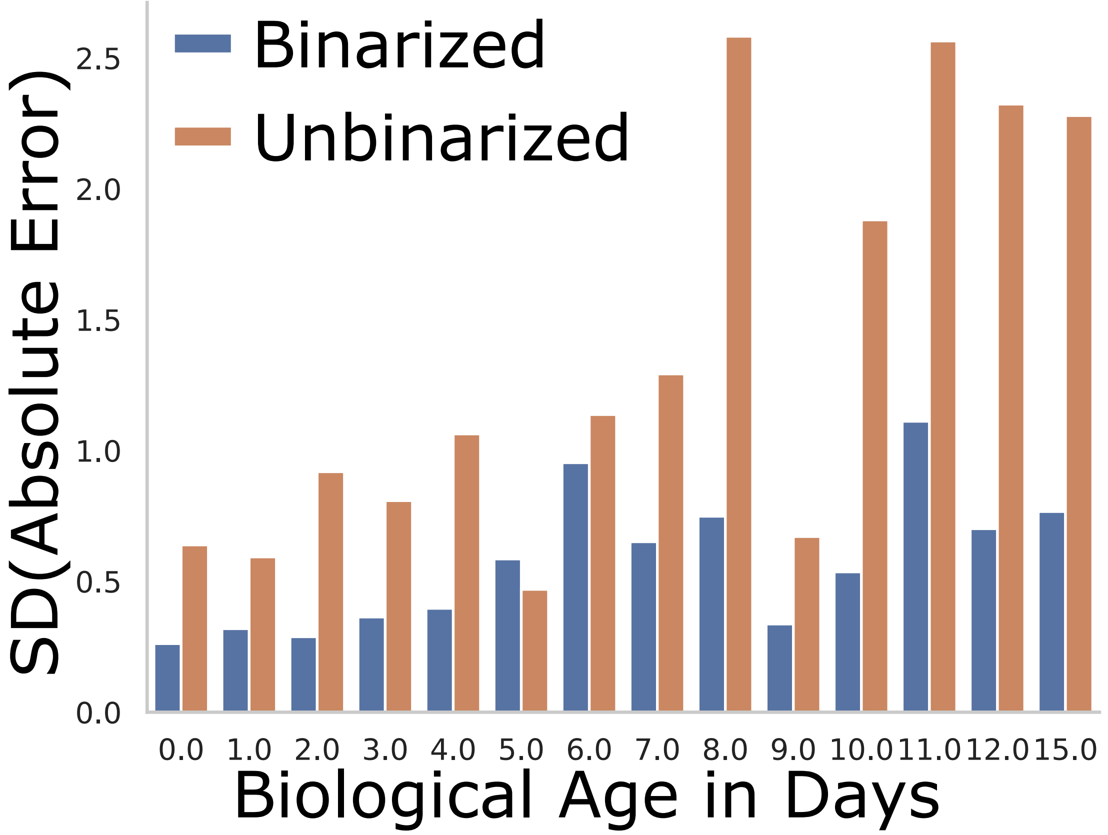

**Fig. S2** **Meyer and Schumacher**

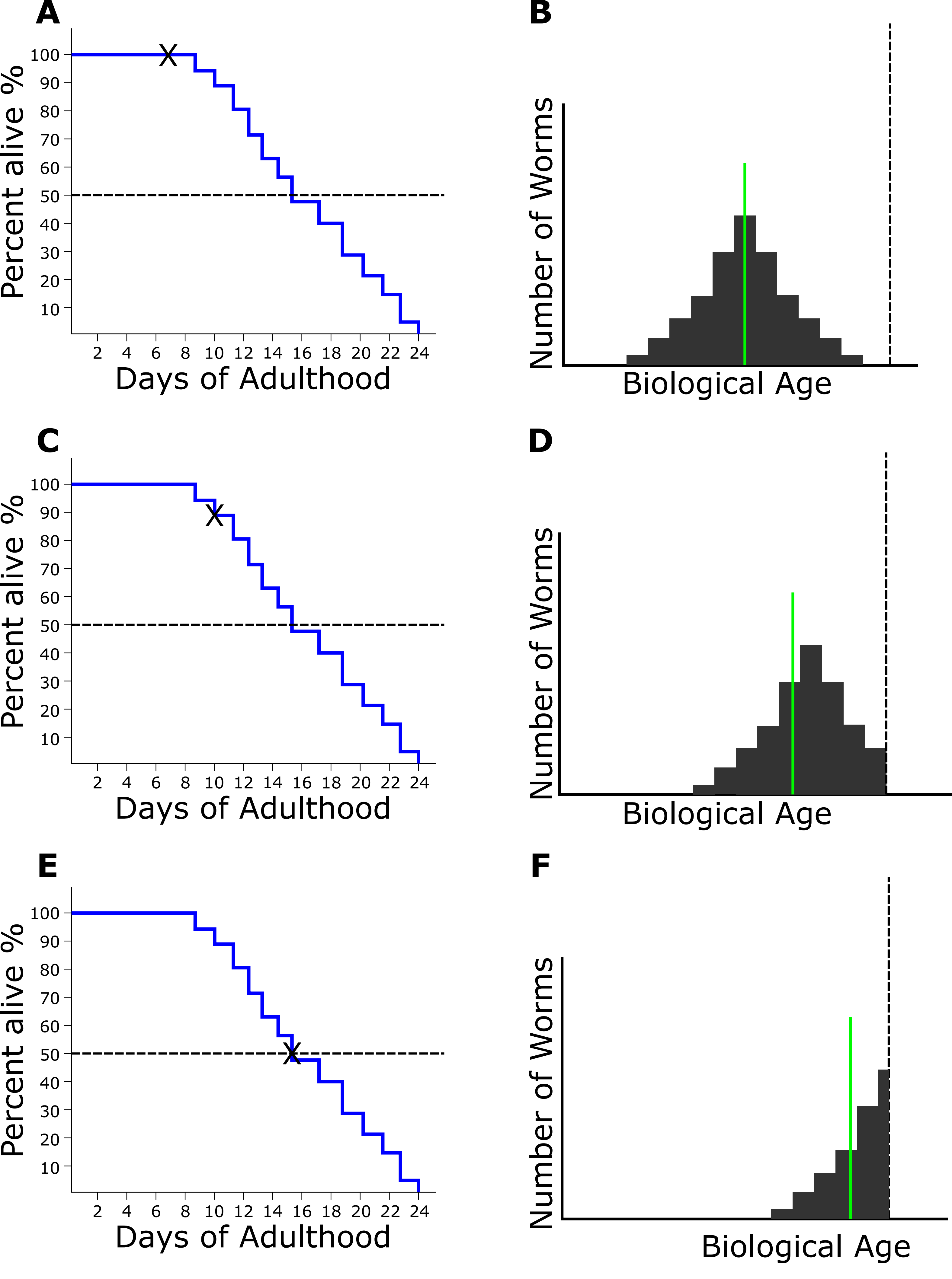

**Fig. S3**

**Meyer and Schumacher**

A

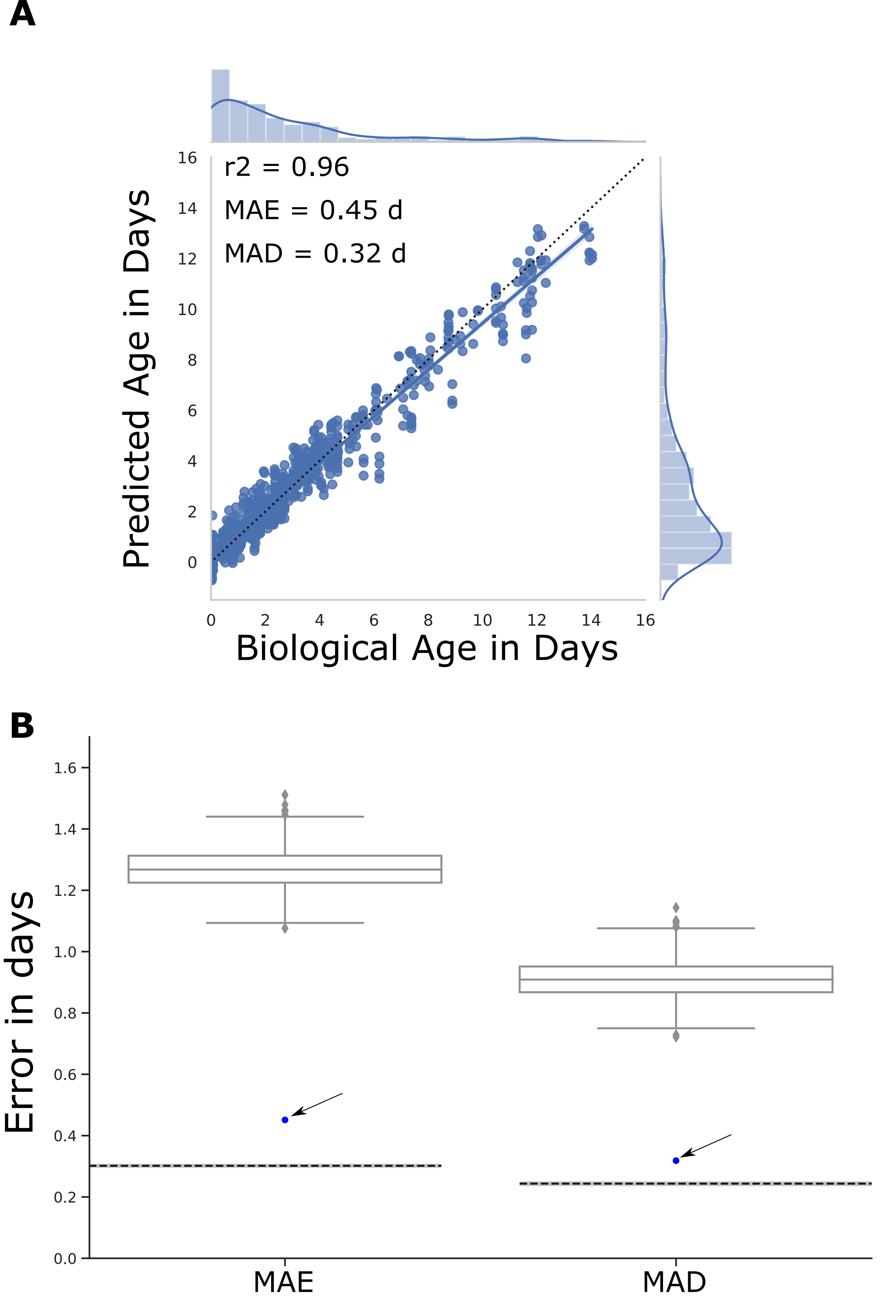

**Fig. S4** **Meyer and Schumacher**

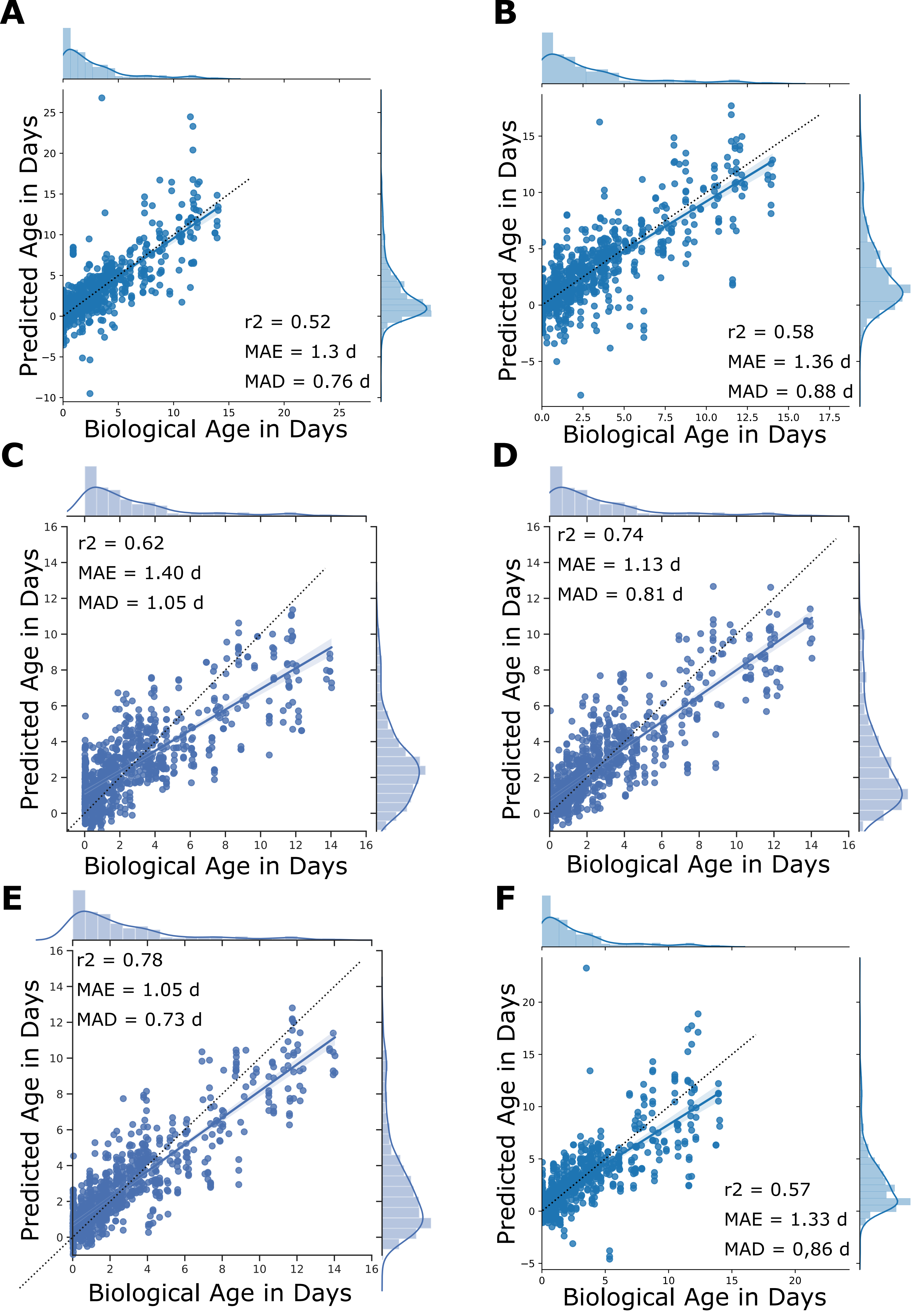

**Fig. S5** **Meyer and Schumacher**

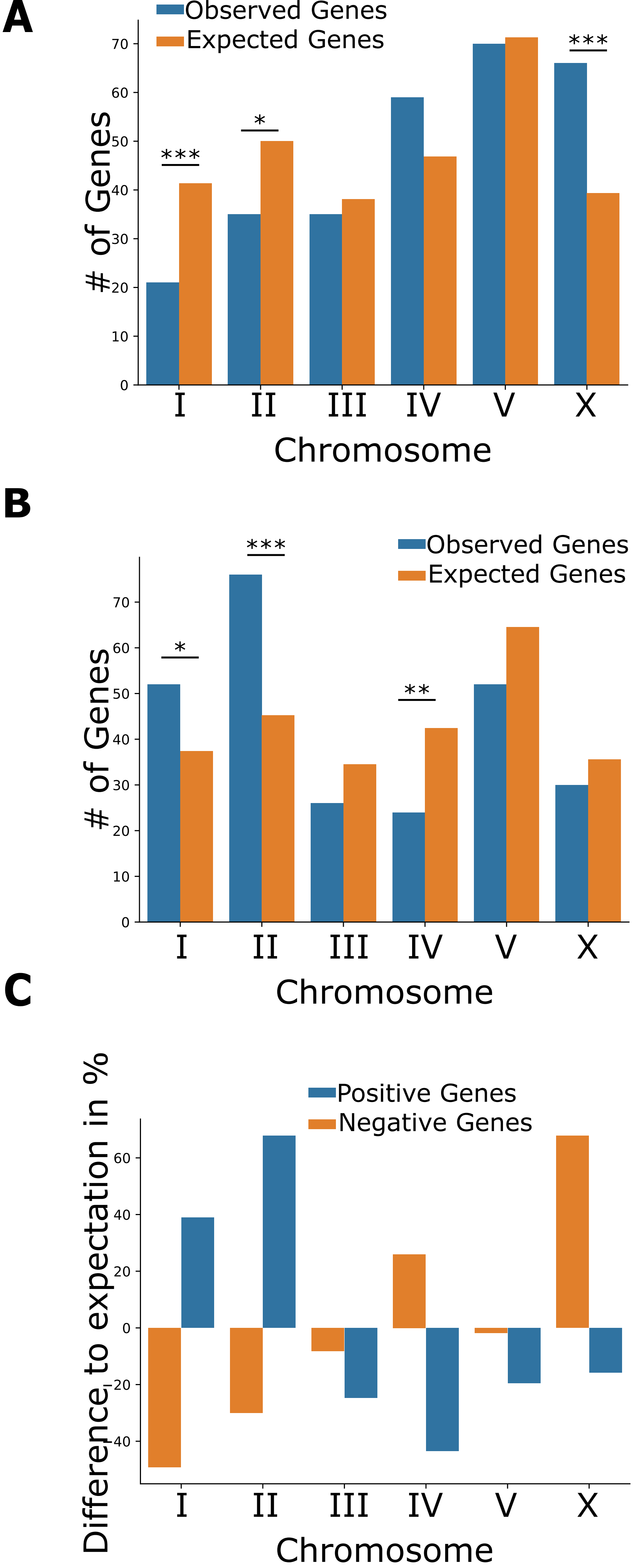

Fig. S6

Meyer and Schumacher

## A

| Name | Motif | % of Genes | % of BG | FC | p-value | q-value |
|---|---|---|---|---|---|---|
| PQM-1(Gata) | 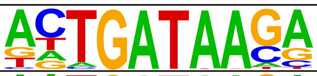 | 29.93 | 17.10 | 1.75 | 1.09E-07 | 1.2E-06 |
| ELT-3(Gata) | 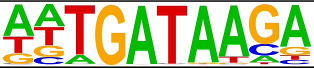 | 31.02 | 20.59 | 1.51 | 3.00E-05 | 1.65E-04 |

## B

| Name | Motif | % of Genes | % of BG | FC | p-value | q-value |
|---|---|---|---|---|---|---|
| ELT-3(Gata) | 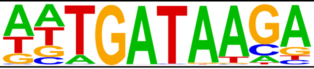 | 30.04 | 19.91 | 1.51 | 7.74E-05 | 8.51E-04 |
| PQM-1(Gata) | 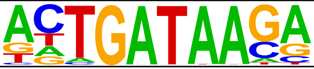 | 23.72 | 16.52 | 1.44 | 1.97E-03 | 0.01 |

**Fig. S7** **Meyer and Schumacher**

**A**

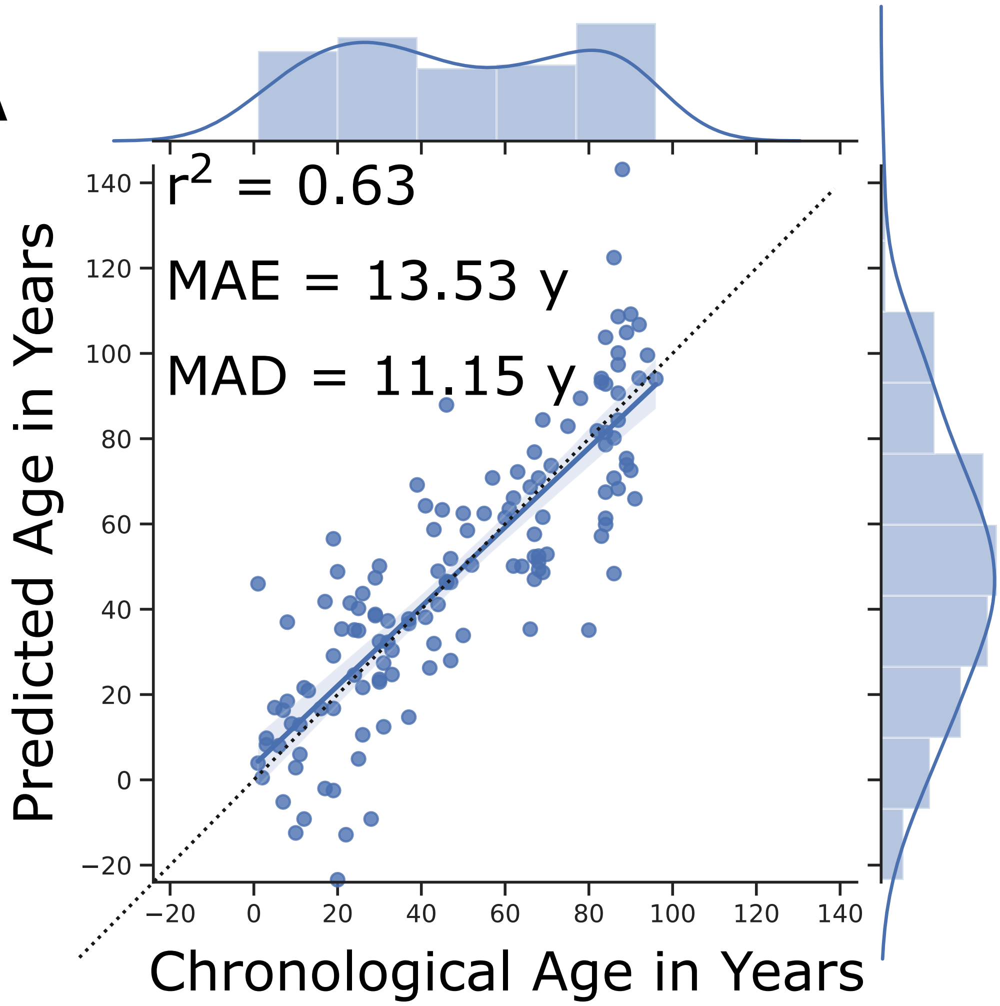

**B**

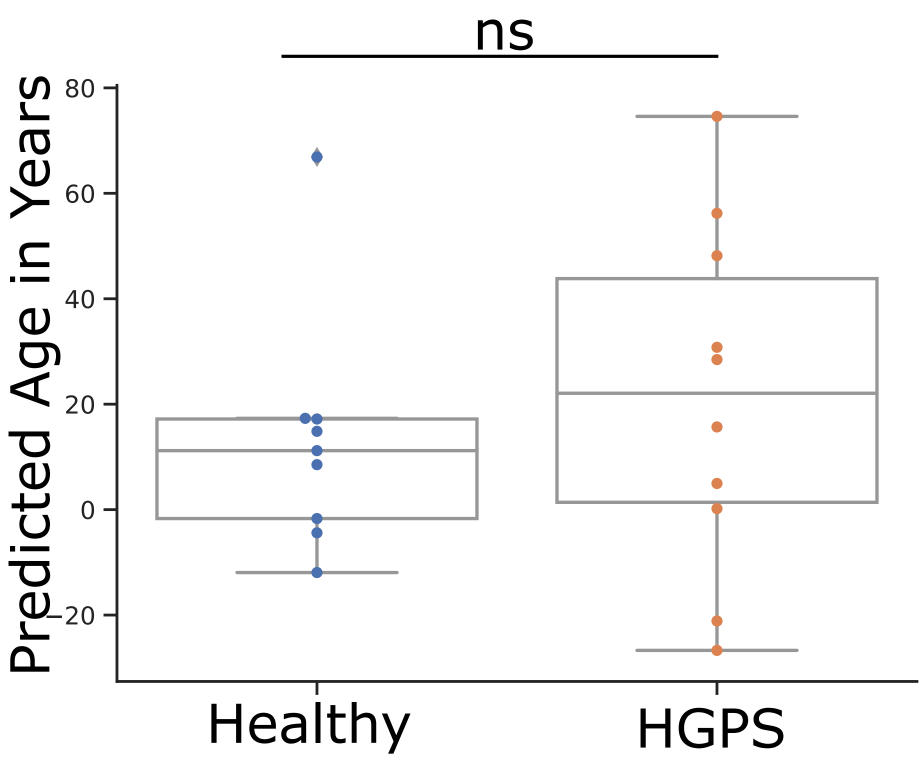

**Fig. S8** **Meyer and Schumacher**
