## Supplementary material for "A transcriptome based aging clock near the theoretical limit of accuracy": FileS1_PythonCode

```
import numpy as np
import pandas as pd
from sklearn.linear_model import ElasticNet
from sklearn.utils.testing import ignore_warnings
from sklearn.exceptions import ConvergenceWarning
import scipy.special


def make_binary(df, filter_genes='WBG'):
    '''

    :param df: Pandas DataFrame with a row for each Sample.
    Columns contain Genes, as well as meta-data,i.e.
    the Strain, Treatment, RNAi, Biological Age, GEO accession number
    :param filter_genes: Compute the binarization only on the genes, mask meta-data
    :return: A binarized copy of the original data with relevant meta-information
    '''
    df_div = df.copy()
    df_div = df_div.filter(regex=filter_genes)
    df_div[df_div == 0] = np.nan
    df_div['Median'] = df_div.median(axis=1)
    df_div = df_div.filter(regex=filter_genes).div(df_div.Median, axis=0)
    df_div[df_div.isna()] = 0
    df_div[df_div <= 1] = 0
    df_div[df_div > 1] = 1

    df_div['Name'] = df.Name
    df_div['Bio_Age'] = df.Bio_Age # biological age in hours of the sample
    df_div['Round_Age'] = np.round(df.Bio_Age / 24) # biological age rounded in days
    df_div['GEO'] = df.GEO
    df_div['Strain'] = df.Strain
    df_div['Treatment'] = df.Treatment
    df_div['RNA_interference'] = df.RNA_interference

    return df_div


def get_xydata(df, y_type='Bio_Age', filter_genes='WBG'):
    '''
    Return the x and y values for the prediction algorithm
    :param df: Pandas DataFrame with a row for each Sample.
    Columns contain Genes, as well as meta-data,i.e.
    the Strain, Treatment, RNAi, Biological Age, GEO accession number
    :param y_type: Age_Perc or Hours
    :param filter_genes: WBG for elegans, ENSG for human
    :return
    '''

    x = df.filter(regex=filter_genes).values
    y = df.reset_index()[y_type].values

    return x, y


def return_subset_of_dataframe(df, subset_col, add_cols=None):
    '''
    Returns the data for a subset of genes
    :param df:Pandas DataFrame with a row for each Sample.
    Columns contain Genes, as well as meta-data,i.e.
    the Strain, Treatment, RNAi, Biological Age, GEO accession number
    :param subset_col: List of genes, i.e. the 576 predictor genes

    add_cols = ['Name', 'Bio_Age', 'Strain', 'Treatment',
    'RNA_interference', 'GEO', 'Round_Age']
    :return:
    '''
    if add_cols:
        for ac in add_cols:
```

```
            if ac not in subset_col:
                subset_col.append(ac)

    return df[subset_col]


def return_regr_fit(x, y, maxiter=1000, alpha=0.075, l1_ratio=0.3, positive=False):
    '''
    Train an Elastic net regression on x and y
    :param x: RNA-seq data for all samples
    :param y: biological age of the samples
    :return:
    '''
    regr = ElasticNet(random_state=0,
                      max_iter=maxiter,
                      alpha=alpha,
                      l1_ratio=l1_ratio,
                      positive=positive)

    regr.fit(x, y)
    return regr


@ignore_warnings(category=ConvergenceWarning)
def cross_validation(df, alpha=0.075, l1_ratio=0.3, age='Bio_Age'):
    '''

    :param df: Pandas DataFrame with a row for each Sample.
    Columns contain Genes, as well as meta-data,i.e.
    the Strain, Treatment, RNAi, Biological Age, GEO accession number
    :param alpha:
    :param l1_ratio:
    :param age:
    :return:
    '''
    cv_tuple = []
    cv_results_dict = {}
    cv_results_dict['Name'] = []
    cv_results_dict['Chrono_Age'] = [] # chronological age of the samples
    cv_results_dict['Bio_Age'] = [] # rescaled biological age
    cv_results_dict['Pred_Age'] = [] # predicted biological age
    cv_results_dict['AE'] = [] # absolute difference between the biological and predicted a
ge
    cv_results_dict['Strain'] = []
    cv_results_dict['GEO'] = []
    cv_results_dict['Treatment'] = []
    cv_results_dict['RNA_interference'] = []

    for i in range(len(df)):
        # only use a samples once
        if (df.iloc[i].Strain, df.iloc[i].Round_Age, df.iloc[i].Treatment,
            df.iloc[i].RNA_interference) in cv_tuple:
            continue
        # set samples aside for testing that have the same strain,
        # rounded biological age, treatment, and RNAi
        test_data = df[
            (df.Strain == df.iloc[i].Strain) & (df.Round_Age == df.iloc[i].Round_Age) &
            (df.Treatment == df.iloc[i].Treatment) & (
                    df.RNA_interference == df.iloc[i].RNA_interference)]
        cv_tuple.append((df.iloc[i].Strain, df.iloc[i].Round_Age, df.iloc[i].Treatment,
                         df.iloc[i].RNA_interference))
        x_test = test_data.filter(regex='WBGene').values
        y_test = test_data.reset_index()[age].values

        # train on everything but the test data
        train_data = df[
            ~((df.Strain == df.iloc[i].Strain) & (df.Round_Age == df.iloc[i].Round_Age) &
              (df.Treatment == df.iloc[i].Treatment) & (
```

```
                   df.RNA_interference == df.iloc[i].RNA_interference))]
        x_train = train_data.filter(regex='WBGene').values
        y_train = train_data.reset_index()[age].values


        trained_model = return_regr_fit(x_train, y_train, alpha=alpha, l1_ratio=l1_ratio)
        predicted_y = trained_model.predict(x_test)

        y_test = list(y_test.flatten())
        predicted_y = list(predicted_y.flatten())

        # compute the absolute errors
        AE = []
        for j in range(len(predicted_y)):
            AE.append(abs(y_test[j] - predicted_y[j]))

        # save the data in the results dictionary
        for h in range(len(test_data)):
            cv_results_dict['Name'].append(
                test_data.iloc[h].Strain + '_' +
                test_data.iloc[h].Treatment + '_' +
                test_data.iloc[h].RNA_interference +
                '_' + str(int(test_data.iloc[h].Bio_Age / 24)))
            cv_results_dict['Strain'].append(test_data.iloc[h].Strain)
            cv_results_dict['Treatment'].append(test_data.iloc[h].Treatment)
            cv_results_dict['GEO'].append(test_data.iloc[h].GEO)
            cv_results_dict['RNA_interference'].append(test_data.iloc[h].RNA_interference)
            cv_results_dict['Chrono_Age'].append(test_data.index[h])
            cv_results_dict['Bio_Age'].append(test_data.iloc[h].Bio_Age)
            cv_results_dict['Pred_Age'].append(predicted_y[h])
            cv_results_dict['AE'].append(AE[h])


    return pd.DataFrame(cv_results_dict)


def correct_Bio_Age(df):
    '''
    Calculate the second correction factor for every sample
    :param df: Pandas DataFrame with a row for each Sample.
    Columns contain Genes, as well as meta-data,i.e.
    the Strain, Treatment, RNAi, Biological Age, GEO accession number
    :return: A copy of df with the updated/corrected biological age
    '''
    df_corrected = df.copy()
    bio = df_corrected.Bio_Age.values
    bio2 = [(y - (scipy.special.erfinv(0.5 - (((scipy.special.erf(((372 - y) / (192 / 3)) /
            np.sqrt(2)) / 2) + 0.5) / 2)) * np.sqrt(2) * 2 * (192/3))) for y in bio]
    df_corrected.Bio_Age = bio2

    return df_corrected
```
